# Neurobiological alterations and autism-like phenotype in mice following regulatory-limit exposure to fungicides in drinking water

**DOI:** 10.64898/2026.10.08.757635

**Authors:** Lucie Salvador-Prince, Anaïs Vignon, Ugo G. Alenda, Pierre-Andre Lafon, Dorine Thobois, Benoit Schneider, Sylvain Lehmann, Carole Crozet, Joan Torrent i Mas, Laurent Chaloin, Michel Vignes, Thibaut Laboute, Jérôme A.J. Becker, Julie Le Merrer, Karine Loulier, Véronique Perrier

**Author notes:** Corresponding authors: Dr. Véronique Perrier, Dr Julie Le Merrer, Dr Karine Loulier.

## Abstract

Neurodevelopmental conditions, including attention deficit hyperactivity disorder (ADHD) and autism spectrum disorders (ASD), have increased markedly over recent decades. This rise parallels the widespread use of agrochemicals, suggesting a contribution of environmental exposures. We hypothesize that pesticide residues may disrupt brain development and promote neurodevelopmental disorders. Here, we test whether concentrations permitted in drinking water affect brain development using a maternal exposure model in wild-type mice. Exposure to a fungicide mixture at 0.3 µg/L, below the current European regulatory limit of 0.5 µg/L, impairs corticogenesis and is associated with alterations in synaptic function and glutamatergic signaling. Offspring exhibit delays in sensory development, reduced social interactions, and increased stereotyped behaviors. We identify the adenosine A2A receptor as a molecular target of anilinopyrimidine fungicides. Together, these findings indicate that low-dose exposure to this fungicide mixture can induce an autism spectrum-like phenotype in the offspring.

## 1. Introduction

Autism spectrum disorder (ASD) is a neurodevelopmental disorder impairing social communications and interactions associated with repetitive behaviors^[1]^. Anxiety, cognitive and motor deficits or epilepsy are comorbidities often associated with ASD^[2]^. The causes of ASD remain unknown, however, twin and familial studies estimated the heritability of ASD to be 50%^[3]^. In addition, epigenetic modifications are detectable in ∼10-20% of individuals with ASD, compared with 1-2% in the general population^[4]^. Heritability studies also suggest that genetic predispositions would be associated with environmental factors, such as heavy metals, medication or pesticides, affecting the mother during her pregnancy^[5]^. Several epidemiological studies revealed that gestational exposure to organophosphate, such as chlorpyrifos, induces memory impairments and decreased IQ performance in 7 years old children^[6]^ and mental deficits in children under 5 years old^[7]^. In addition, cohorts of children exposed to organochloride pesticides are associated with increased ASD risk and intellectual deficiencies^[8]^. Besides, the risk to develop ASD in child is increased when mother’s residence during pregnancy is within 2,000 m of farmlands^[9]^.

Because glyphosate is the most used herbicide in the world, it was suspected to be involved in the recent increase of ASD prevalence worldwide^[10]^. Indeed, epidemiological data demonstrate a positive correlation between the rise of glyphosate usage on corn and crops in the United States over the years 1995 to 2010 and the increase in ASD incidence rates over the same period^[10]^. However, epidemiological studies are retrospective and cannot determine precisely which chemicals have the greatest potential to alter the neurodevelopment, and by which cellular and molecular mechanisms they induce ASD phenotype. Thus, animal models chronically exposed to this herbicide were developed to identify a possible causal link with ASD phenotypes. Recent study showed that pre- and post-natal exposure to high levels of glyphosate administered through water (at 980 mg/L corresponding to a 158 mg/kg b.w./day) induced autism-like behaviors in offspring^[11]^. Nevertheless, the acute dose used did not reflect daily human exposure. Glyphosate resistant weeds have begun to develop around the world and the controversy regarding the glyphosate ban in Europe has prompted phytosanitary industry to replace glyphosate by glufosinate ammonium (GLA). Interestingly, a study by Laugeray et al. (2014)^[16]^ showed that pre- and post-natal administration of GLA at 1 mg/kg (dose of 30 µg, 3 times a week), in female mice, induced autism-like phenotypes in offspring. This intranasal administration of GLA mimics aerosol exposure to pesticides and doses administered are particularly relevant to agricultural communities including farmers that are highly exposed to pesticides by air contamination. Concerning the general population, although 30 µg seem low, this concentration is high compared to pesticide levels measured in the urban air, ranging between 0.2 and 1 ng/m^3^, corresponding to 2.8-14 ng/day/person given that an individ ual breathes 14 m^3^ of air/day^[13]^. This model suggests that a part of ASD cases could thus be related to the exposure of populations to pesticide-contaminated air. However, air contamination alone would not explain the frequency of ASD in the general population. Other routes of contaminations such as water and food may play a role in the rising prevalence of ASD, as well as other pollutants.

Here, we studied the contribution of pesticides present in water by testing the regulatory limit dose allowed for human consumption, 0.1 µg/L for one pesticide or 0.5 µg/L for pesticide cocktail. According to the European Union (EU) directives on water^[14]^, these low doses of pesticides have been selected to « be consumed safely on a lifelong basis and thus represent a high level of health protection ». We focused on anilinopyrimidine fungicides (cyprodinil, mepanipyrim and pyrimethanil) which are widely used to preserve crops and for the long-term storage of fruits and vegetables. Many reports have established the presence of anilinopyrimidine residues in our environment (air^[13]^, water^[15]^ and many food products^[16]^). More worrying, these three fungicides were even detected in baby food jars^[17]^ and recent epidemiological studies revealed the presence of anilinopyrimidine residues in hair and blood samples of pregnant women in ELFE and ISA cohort respectively^[18],[19]^. Previously, we developed a model of contamination from mother to offspring by adding 0.1 µg/L of cyprodinil, mepanipyrim or pyrimethanil alone, or 0.3 µg/L in combination, in drinking water below the regulatory limit dose for pesticide mixture (<0.5 µg/L)^[20]^. We already showed that this gestational exposure corrupted neurogenesis in neonates as early as post-natal day 3 (P3). Besides, properties of adult neural stem cell isolated from 3 months old exposed-animals, were impaired especially proliferation, differentiation, as well as migration^[20]^.

Our working hypothesis is that neural stem cells affected by pollutants during brain development could induce defects in cortical development through alterations of neuronal and glial cells properties, impairments in neuronal network organization and synaptic transmiss io n. Such defects may contribute to altered behavioral outcomes and increase susceptibility to neurodevelopmental disorders such as ASD. To test our hypothesis, we exposed wild-type (WT) female mice to a cocktail of the 3 anilinopyrimidine fungicides in water, as previously reported^[20]^, to study the impact on the neurodevelopment of their offspring chronically exposed from gestation to culling. In utero electroporation using MAGIC markers fluorescent reporter transgenes^[21]^ allowed us to identified several defects in the cerebral cortex of exposed offspring, including alterations of the morphology and spatial distribution of pyramidal neurons. Electrophysiological studies performed on hippocampal slices from cocktail exposed neonates revealed polysynaptic neuronal signals reminiscent of seizure-like activity and impairments of synaptic plasticity. These defects are associated with phenotypic developmental delays, and later, with social interaction and social novelty deficits associated with stereotypies in adult mice. Biochemical studies showed a blockade of adenosine (A2A) receptors by these fungicides. Altogether, these data suggest that pre- and post-natal exposure to a cocktail of fungicides at regulatory limit dose via tap water lead to a chemo-induced model of ASD in mice, while identifying a mechanism of action underlying these environmental chemica l-induced developmental defects.

## 2. Results

### 2.1 Gestational exposure to fungicide cocktail induced neuroanatomical defects in the cerebral cortex of P10 neonates

To evaluate the impact of chronic exposure to low doses of fungicides on cortical development, we performed *in utero* electroporation (IUE)-based MAGIC Markers (MM) strategy in E13.5 embryos of pregnant C57Bl/6J females exposed or not to the cocktail of anilinopyrimid ine fungicides (Figure 2a). This multicolor reporter approach enables to mark cortical progenitors and to track their entire neuronal and glial descent ^[22]^. Following PiggyBac (PB) transposase-based genomic integration and Cre-recombination of the PB-CAG-Cytbow transgenes, cyan, yellow, or red fluorescent proteins (FP) are expressed, yielding to a semi sparse mosaic of color combinations in cortical progenitors, pyramidal neurons and astrocytes. This multicolor fate mapping strategy grants access to cell morphology and location in upper and deep cortical layers (Figure 2). Neuroanatomical analyses revealed substantial deformations of different cortical layers in several brains of cocktail-exposed animals at 10 days post-natal (P10) that were not found in CTR animals (Figure 2b). We also observed cytological abnormalities as cortical mantle cysts (Figure 2c, cocktail 2-3), corresponding to clusters of deep-layer pyramidal CTIP2^+^ neurons aggregated at the surface of the cortex in cocktail exposed animals compared to CTR, reminiscent of neuronal migration defects^[23]^. These cortical cysts present similarities to cobblestones present at the surface of the cerebral cortex and found in human pathologies such as focal cortical dysplasia and lissencephalies. An increase in astrocyte cluster size, as well as a higher number of astrocytes per cluster, was observed in exposed animals compared to controls (Figure 2d, CTR vs Cocktail, enlargements 1–2 vs 3–4). However, these changes were observed qualitatively but could not be precisely quantified due to limitations of the electroporation procedure, as the number of electroporated cells varies between animals. Furthermore, the software was technically limited in its ability to detect all astrocytes present within the cluster. Within upper layers, we noticed an abnormal spatial distribution of pyramidal neurons displaying aggregated cell bodies trapped in interlacing fibers, instead of a well-organized structure of parallel dendritic fibers (Figure 2e). We also observed that numerous CTIP2^+^ neurons presented an abnormal “multibranched” morphology, reminiscent of immature multipolar neurons, in cocktail-exposed animals (Figure 3a). Neuronal reconstruction using Vaa3D^[24]^ revealed increased branching (ns), node number (*p*= 0.0116), and total volume (*p*= 0.0093) in cocktail-exposed animals compared to controls (Figure 3b).

**Figure 1.**
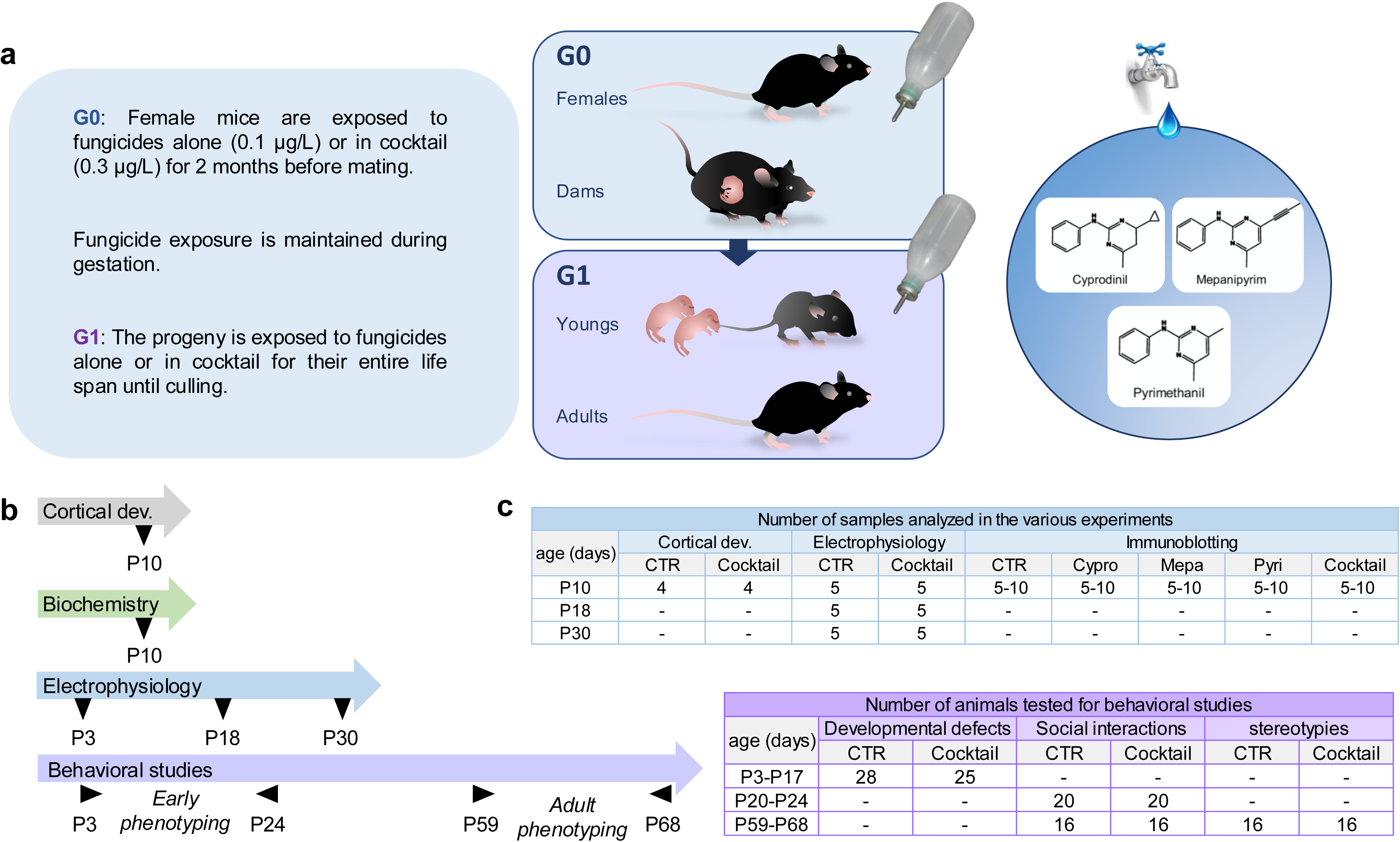
Study design illustrating the protocol of exposure of WT female mice to fungicide cocktail, the type of experiments conducted, and mice numbers. (a) Female mice (G0) were exposed to fungicide, alone or in cocktail, from 2 months before mating, until weaning of their pups. Newborns (G1) were continuously exposed to the fungicides from *in utero* to adult stages. Chemical structures of the 3 anilinopyrimidine fungicides: cyprodinil (cypro), mepanipyrim (mepa) and pyrimethanil (pyri). The dose of fungicide used in water was 0.1 µg/L each. This dose corresponds to the regulatory limit dose allowed by EU in water for human consumption (EU council directive 98/83EC). (b) Multiple experiments were performed to characterize the impact of fungicides, alone or in cocktail on mice, including neuroanatomical studies performed at P10 following *in utero* electroporation of MAGIC Markers, biochemistry (P10), electrophysiological recordings on acute brain slices (P10-P30) and behavioral evaluations in young (P3-P24) and adult mice (2 months). (c) Tables summarizing the number of mice for each type of analysis.

**Figure 2.**
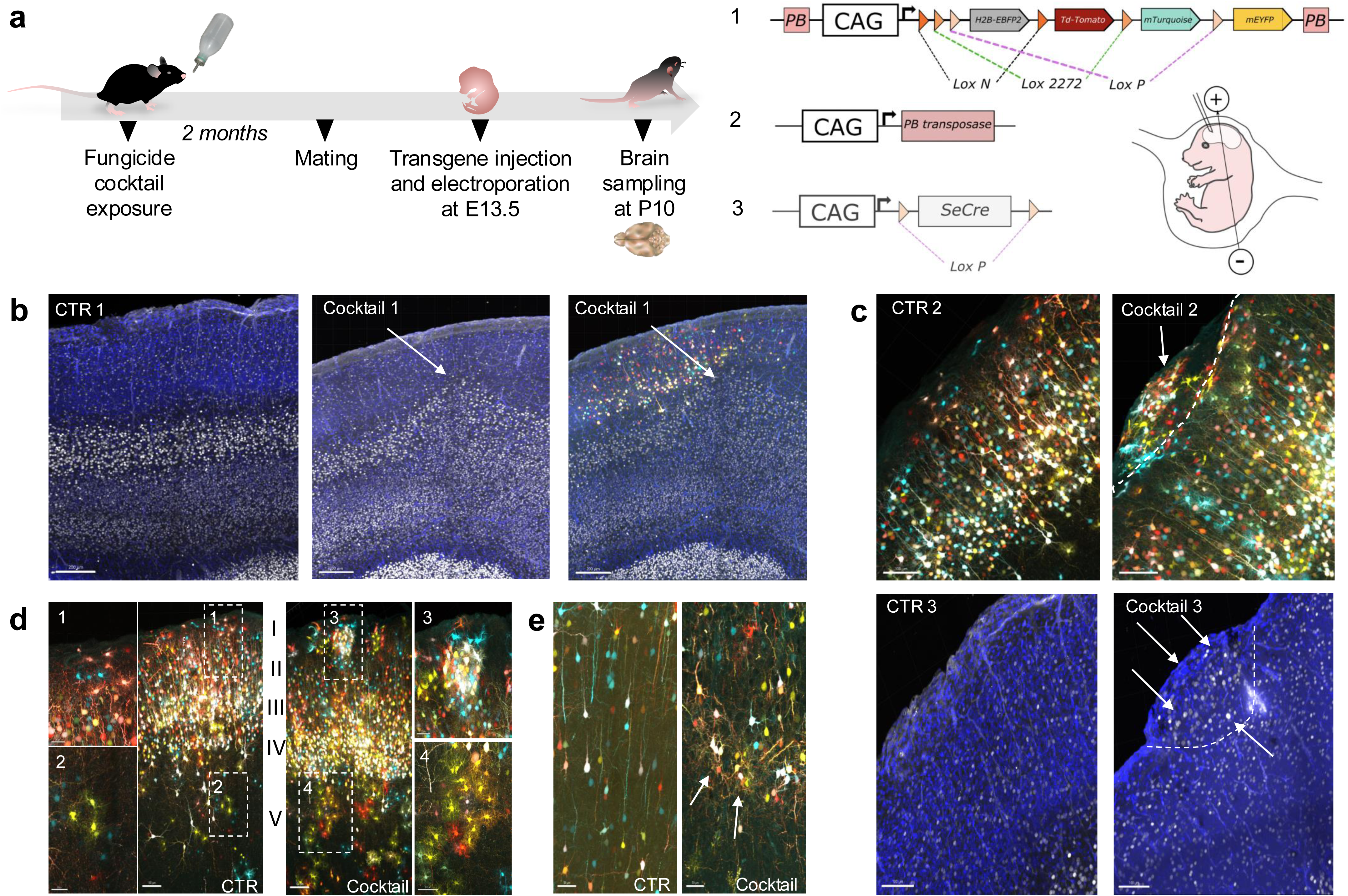
*In utero* electroporation with MAGIC marker plasmids reveals various abnormalities in cortical layers in cocktail-exposed mice. (**a**) Experimental protocol used for the *in utero* electroporation of mouse embryos at E13.5 exposed or not to the cocktail of fungicides and description of the electroporated plasmids: (1) PB-CAG-Cytbow, (2) PiggyBac transposase, and (3) SeCre. (**b**) Immunofluorescence of brain tissue sections of CTR (CTR1) mouse, or cocktail-exposed mouse (Cocktail1) at P10. Illustration of cortical layer deformatio ns labeled with CTIP2, a specific marker of layers V-VI (Cocktail1), as well as deformations of upper cortical layers showing abnormal positioning of electroporated cells (right image) at the surface of the cerebral cortex in P10 mouse exposed to the cocktail of fungicides. The blue labeling corresponds to Hoechst cell nuclei staining. (**c**) Confocal images of 2 different CTR (CTR2 and CTR3) and cocktail-exposed mice (Cocktail 2 and 3) showing abnormalities of the upper cortical layer, with mantle cysts, delineated by dotted lines. (**d**) Representative images of cerebral cortices from P10 animals electroporated at E13.5 with the MAGIC Markers constructs, showing large clusters of electroporated multicolor cells in cocktail-exposed mouse (dotted frame 3, and enlargement 3) which are absent in the CTR mouse (dotted frame 1 and enlargement 1). Large astrocyte clusters in the lower cortical layers are also visible in a cocktail-exposed mouse (dotted frame 4 and corresponding enlargement), absent in the CTR (dotted frame 2 and corresponding enlargement) animals. (**e**) Representative images of the disorganization of pyramidal neurons within the cortical layers, observed in mice exposed to the cocktail of fungicides (right image) compared to an equivalent area in CTR mice (left image) with the white arrows pointing to the reported abnormalities.

**Figure 3.**
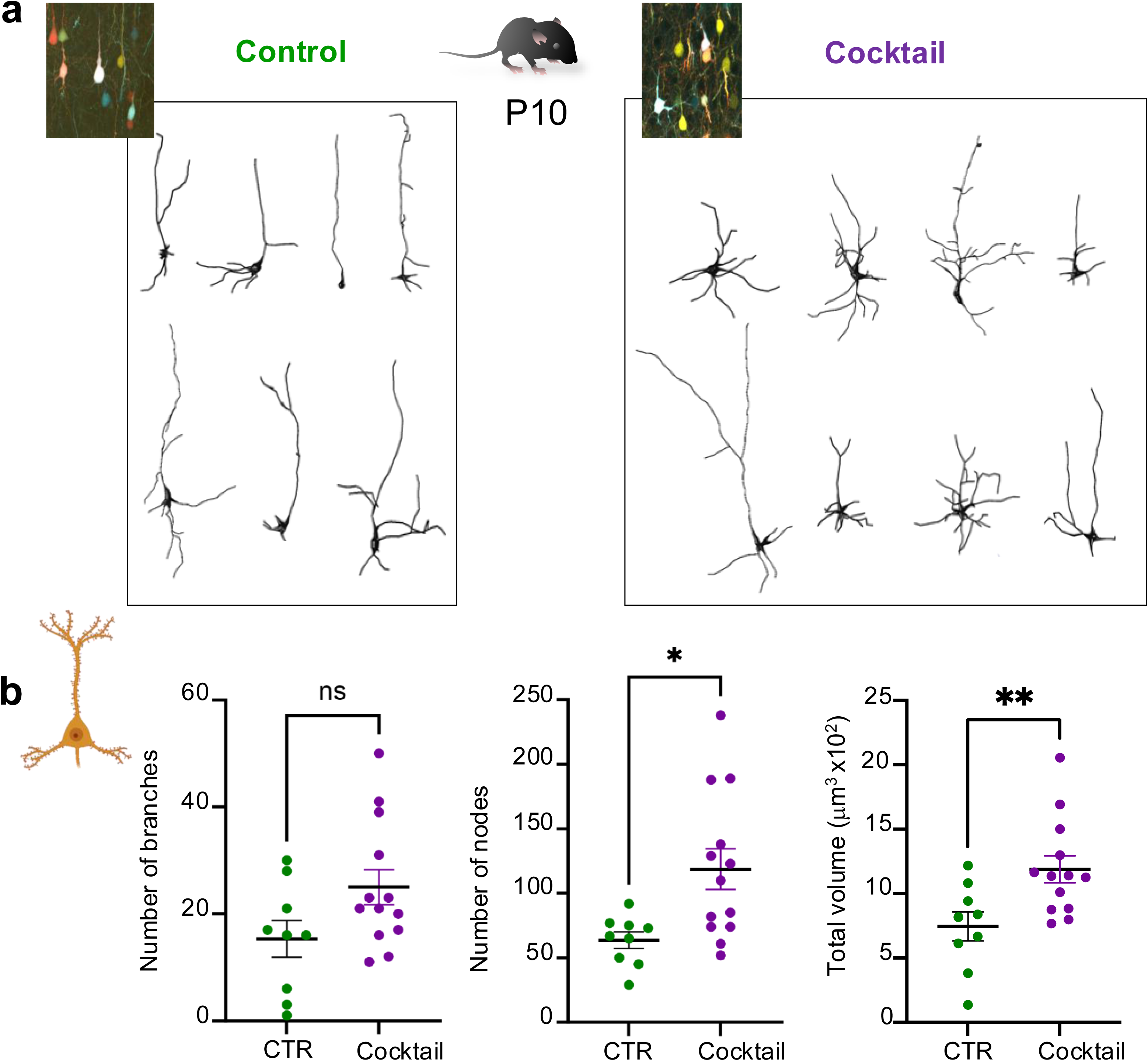
In utero exposure to fungicide cocktail results in multibranched pyramidal neurons in the cortex of fungicides exposed-mouse. (**a**) Representative 3D reconstructions of cortical pyramidal neurons in control (CTR) and cocktail-exposed mice (Cocktail) at P10. Abnormal morphologies, as multibranched neurons are reported in fungicides-exposed mice (CTR n=4, Cocktail n=5). (**b**) Numbers of branches, nodes and total volume were quantified from re-constructed neurons (CTR n=9, Cocktail n=13) issued from different animals (CTR n=4, Cocktail n=5). Number of nodes: unpaired T-test, *p*=0.0116. Total volume of neurons: unpaired T-test with Welch’s correction, *p*=0.0093. The 3D reconstructions and quantificat io ns of neurons were done using VAA3D software and the associated plugin neuron tracing “APP2” from Peng lab (Columbia University, NY).

### 2.2 Gestational exposure to fungicides impaired immature and differentiated neuron populations and increased astrocytes through the PI3K/Akt and Wnt/β-catenin pathways

MAGIC Markers multicolor reporter strategy has underlined several neuroanatomical defects in the cerebral cortex of animals exposed to fungicide cocktail, as well as clusters of astrocytes and pyramidal neurons suggesting alterations in neuronal and glial production. We thus perform biochemical analyses to evaluate if the *in utero* exposure to the cocktail of fungicides may impair the proportion of neural cell populations.

First, we quantified by western blot the expression levels of DCX, NeuN and Aldh1l1 expressed by immature neurons, mature neurons and astrocytes respectively (Supplementary Figure S1). At P10 days, DCX expression is strikingly decreased by 50%, in cocktail-exposed animals compared to CTR group (*p=*0.027), as well as NeuN^+^ (cocktail: *p=*0.02) (Figure 4a-b). By contrast, Aldh1l1 expression was increased in the cocktail group (p = 0.002; Figure 4c), supporting enhanced astrocyte production, consistent with the numerous astrocytic clusters revealed by MAGIC Markers lineage-tracing experiments (Figure 2d). These results indicate modifications in the proportion of immature and differentiated neural cell populations upon fungicide exposure, suggesting some alterations in the production of neurons and glial cells. Thus, we analyzed cell signaling pathways known to be involved in cell differentiation and proliferation during brain development. Previous studies showed following its phosphoryla t io n on S552 (P-β-cat), the β-catenin protein is translocated into the nucleus and regulates the activation of genes involved in the self-renewal of neural progenitor cells^[25]^ and neuronal differentiation^[26]^ during postnatal neurogenesis (Figure 4d). We determined in our samples whether β-cat could be involved in this process. Analysis on P10 neonate’s brain homogenates showed a 2-fold increased levels of the ratio P-Akt (S473)/Total Akt, upon cocktail exposure (cocktail: *p* < 0.0001) (Figure 4e, Supplementary Figure S1). Analyses by western blotting of P-β-cat (S552)/Total β-cat also revealed a 2-fold increase of the ratio upon cocktail exposure compared to CTR group (cocktail: *p=* 0.002) (Figure 4f, Supplementary Figure S1). These data suggested that the modifications of expression of neuronal (DCX^+^, NeuN^+^) and astrocytic (ALDH1L1^+^) markers could be linked to P-β-cat (S552) accumulation, through the cross-talk of PI3K/Akt and Wnt/β-catenin signaling pathways.

**Figure 4.**
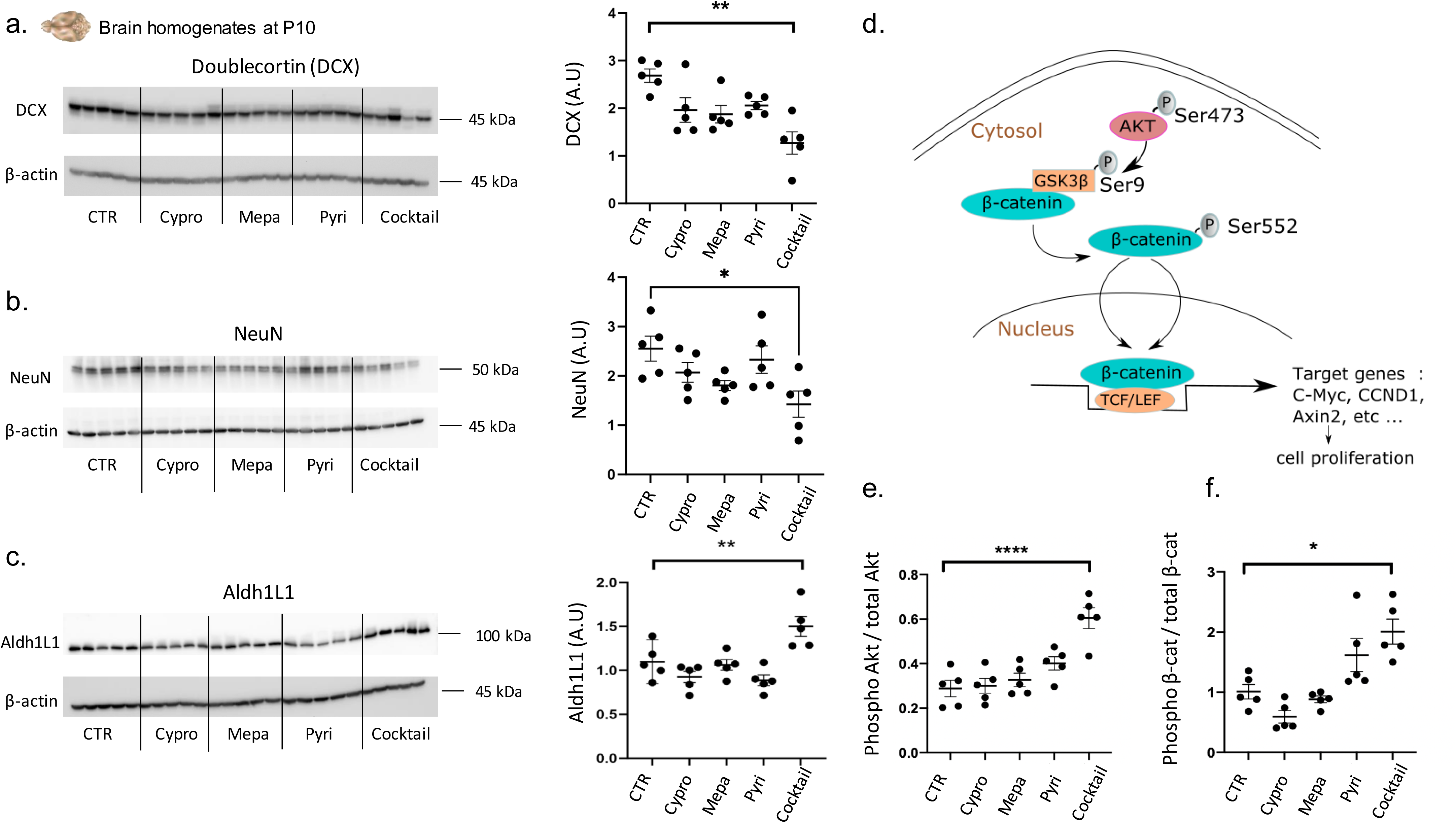
Involvement of the PI3K/Akt and Wnt/β-catenin signaling pathways in modifying the expression of various neural population markers. The expression of DCX (immature neurons) (**a**), NeuN (mature neurons) (**b**), and ALDH1L1 (astrocytes) (**c**) proteins was analyzed in brain homogenates of newborns at P10 exposed during gestation to fungicides, alone or in a cocktail (0.1μg/L per fungicide) or with 3x10^-5^ % DMSO (CTR). Protein loading control was performed with β-actin. (**d**) Graphical representation of the PI3K/Akt and Wnt/β-catenin crosstalk. The phosphorylation of Akt on its Ser473 will allow the phosphorylation of GSK3β on its Ser9 site, thus inhibiting the degradation of β-catenin. The latter will then accumulate in the cytosol. Once phosphorylated on its Ser552 site, β-catenin will be able to translocate to the nucleus and associate with TCF/LEF transcription factors and thus activate a whole family of genes, some of which are involved in cell proliferation. The amount of Phospho-Akt (Ser473)/Total-Ak (**e**) and Phospho-β-catenin (Ser552)/total-β-catenin (**f**) were assessed by immunoblotting. β-actin was used as a loading control. Note: AU, arbitrary units; CTR, control; DMSO, dimethyl sulfoxide; SEM, mean standard error, n = number of samples, p, p-value.

### 2.3 Gestational exposure to fungicides modified postsynaptic proteins and induce d epilepsy-like phenomenon in brain tissue sections

Activation of the β-catenin signaling pathway has been shown to increase dendritic arborization in hippocampal neurons^[27]^ and regulate post-synaptic proteins such as PSD95 and NMDA receptors^[28]^. Thus, we wondered if the increased levels of P-β-cat (S552) in P10 neonates upon exposure to fungicides could impact proteins involved in the glutamatergic post-synaptic transmission (Figure 5a, Supplementary Figure S2). Immunoblotting revealed an increase of PSD95 levels by about 2-folds (mepa: *p=* 0.008) in P10 mice exposed to fungicides alone (Figure 5b, Supplementary Figure S2), as well as a 2-fold increase of Phospho-CAMKII/total CAMKII levels (mepa: *p=*0.009; pyri: *p=*0.016) compared to their control littermates.

**Figure 5.**
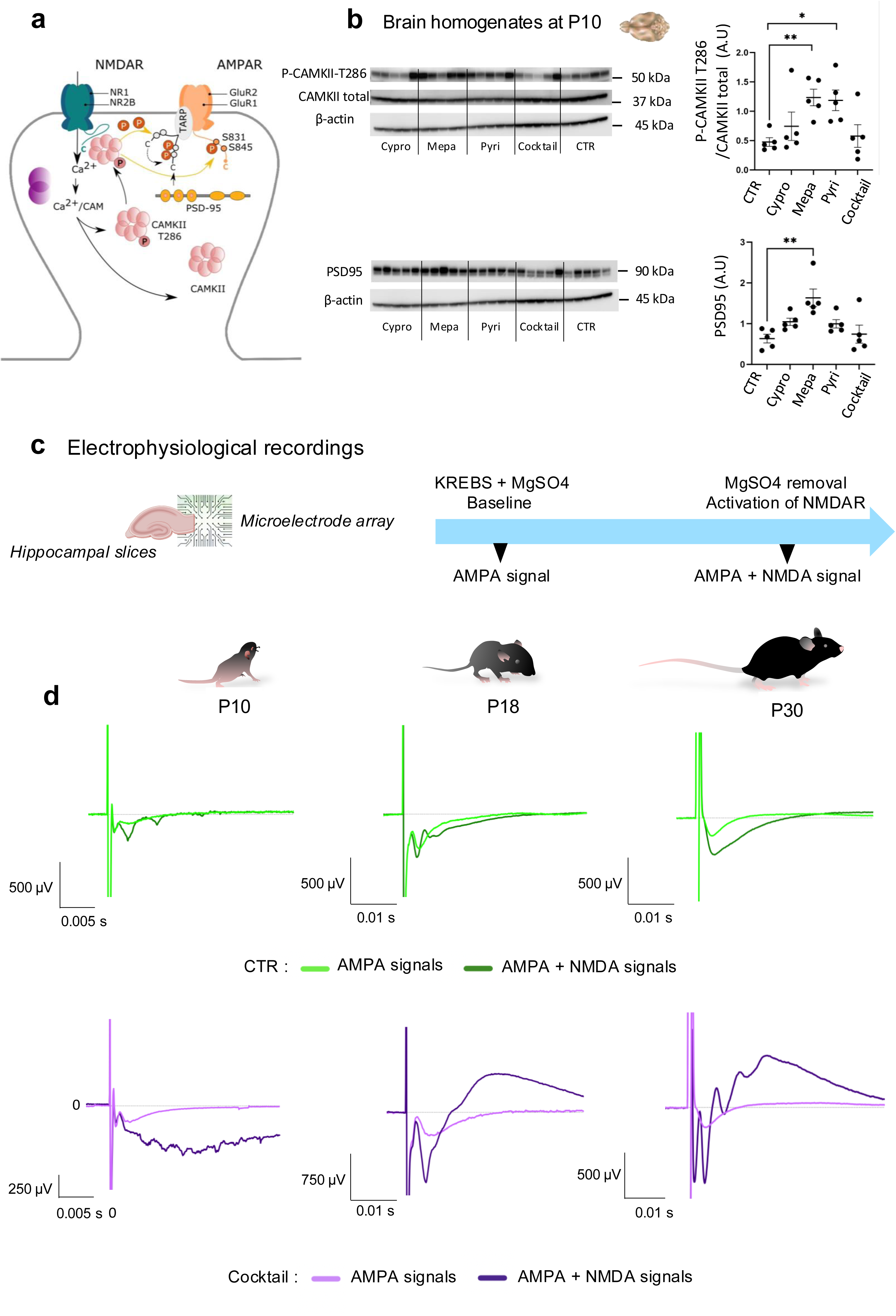
Exposure to fungicides during gestation and lactation alters post-synaptic protein expression and induces epilepsy-like events. (**a**) Representative schema of the different actors of post-synaptic glutamatergic transmis s io n following calcium entry via GluN2B subunit-containing NMDA receptor. Protein expression levels of (**b**) P-CAMKII-T286/total-CAMKII ratio and PSD95 and their quantifications. All signal intensities were normalized from β-actin signal. For the P-CAMKII-T286/total CAMKII ratio, CTR vs Mepa: p=0.0049, vs Pyri: p=0.0064. n = 5 different mouse brains per group (CTR or cocktail). For PSD95, CTR vs Mepa: p=0.0209, vs Pyri: p=0.0010. The displayed P-CAMKII, total CAMKII and PSD95 labelings are from the same membrane that has been stripped and therefore share the same control (β-actin). (**c**) Electrophysiology protocol applied to hippocampal slices from animals exposed or not to the cocktail of fungicides during gestation and lactation; (**d**) Up, representative AMPA receptor dependent fEPSPs (light green curve) and AMPA and NMDA receptors dependent fEPSPs (dark green curve) recorded in Mg^2+^ ions-free solution from CTR mice. Down, illustrative AMPA receptor dependent fEPSPs (light purple curve) and AMPA and NMDA receptors dependent fEPSPs (dark purple curve) recorded in Mg^2+^ ions-free solution from cocktail-exposed mice. Each curve is an average curve of a 2-min recording when the signal is stable, before or after Mg^2+^ ions removal from extracellular medium. Recordings were performed in hippocampal acute slices from animals at P10, P18 or P30.

Since we showed alterations in the expression levels of post-synaptic proteins, we investigated next whether such modifications could have consequences on synaptic transmission and plasticity. Indeed, changes in PSD95 and CamKII expression by fungicides exposure could change NMDA receptors availability and responsiveness and thus impact on synaptic plasticity^[29]^. Thus, electrophysiological recordings were undertaken on acute hippocampal slices from animals exposed to fungicide cocktail during gestation (P10, P18 or P30) with Multi-Electrode arrays (Figure 5c-d). We examined the contributions of AMPA and NMDA receptors to basal excitatory post-synaptic potentials (Figure 5d). For this, recordings were first performed in the dendritic field of CA1 hippocampal neurons in the presence of Mg^2+^ ions in the extracellular medium in order to record AMPA receptor dependent fEPSP (Figure 5d: light green and light purple). Mg^2+^ ions were then removed from the extracellular medium in order to reveal NMDA dependent synaptic signals (fEPSP_AMPA + NMDA_) in addition to the AMPA signals (Figure 5d: dark green and dark purple). In hippocampal slices from animals exposed to fungicides *in utero* (Cocktail), fEPSP recordings showed drastic alterations right after Mg^2+^ ions removal: fEPSP were strikingly enhanced from P10 and exhibited multiple spikes at later stages (Figure 5d) previously described as epileptiform field potentials (EFPs). Those abnormalities were frequently detected in fungicide exposed animals (between 40 and 80%) used for electrophysiology experiments (Supplementary Table S4).

### 2.4 Delay of the GluN2B/GluN2A switch in NMDA receptors in neonates exposed to fungicide cocktail

Because epilepsy-like phenomenon was frequently detected in animals exposed *in utero* to fungicides, we analyzed the GluN2B/GluN2A subunit composition of the NMDA receptor (also known as NR2B/NR2A). Indeed, studies have demonstrated the role of GluN2B subunits in epilepsy through neuronal excitotoxicity due to excessive intracellular calcium entrance^[30]^. At birth, synaptic NMDA receptors in the mouse hippocampus are predominantly composed of GluN2B-subunits at the post-synapse membrane^[31]^. During the first post-natal week, many new synapses are formed, however they do not contain GluN2B subunits, making GluN2A the main subunit starting from P7 stage. This modification is named the GluN2B/GluN2A switch (also known as “NR2B/NR2A switch”)^[32]^. Physiologically, this “switch” contributes to the regulation of the critical period of developmental plasticity^[33]^. We thus suspected impairments in the switch process upon fungicides exposure. In order to isolate the NMDA component of the fEPSPs, AMPA receptors were blocked by addition of 10 µM DNQX in the extracellular medium while NMDA were still under Mg^2+^ ions deprivation (Figure 6a-b). To verify this hypothesis, we used ifenprodil, an antagonist preferentially targeting NMDA receptors containing GluN2B subunits. Our result showed that in P10, P18 and P30 control animals (green curves), the addition of ifenprodil did not modified fEPSP_NMDA_ amplitude illustrating the absence of GluN2B subunits at the post-synaptic membrane meaning that the switch already occurred as previously described^[32]^ (Figure 6b). By contrast, in P10 and P18 aged animals that were exposed *in utero* to fungicide cocktail (purple curves), the addition of 10 µM ifenprodil partially inhibited fEPSP_NMDA_ demonstrating the persistence of NMDA receptors containing GluN2B subunit at post-synaptic sites (Figure 6b). However, in P30 animals exposed to cocktail, ifenprodil had no effect, likely due to the disappearance of GluN2B subunits at the post-synaptic membrane, suggesting that the GluN2B/GluN2A switch occurred between P18 and P30 in the cocktail group (Figure 6b). The quantification of signal blocked by ifenprodil showed that for mice previously exposed to the fungicide cocktail, around 25 % of fEPSP_NMDA_ are still due to GluN2B subunit at P10 and P18 (P10: *p=*0.0094; P18: *p=*0.0107) (Figure 6b). As expected, ifenprodil had no effect on fEPSP_NMDA_ in control mice at P10 and P18. Since GluN2B/GluN2A switch is impaired during the critical window of synaptic plasticity development and maturation, we suspected that fungicide exposure could have an impact on NMDA receptor dependent long-term synaptic changes in the hippocampus^65,66,68,70^. We therefore focused on long-term depression (LTD), which is more easily induced during early developmental stages, a period characterized by a higher contribution of GluN2B-containing NMDA receptors. The application of low-frequency stimulation (1 Hz, 15 min) resulted in a 40% reduction in the evoked synaptic response (n = 5) in the CA1 area of CTR animals (Figure 6c). By contrast, LTD was not observed in slices from fungicide-exposed mice (n = 5), as synaptic transmission returned to baseline within 40 minutes after low frequency applicatio n. The difference between the two groups was highly significant (*p*<0.0001) (Figure 6c). Because LTD is a fundamental mechanism for the acquisition of memory processes and learning, the impairments observed suggested potential alterations of these features^[37]^. Thus, groups of adult animals (3 months) were tested for their learning and memory abilities, using the novel object recognition test (NOR). Results showed that animals exposed to the cocktail of fungicides, from in utero up to 3 months, have less interest to explore the novel object, compared to control group (Figure 6d). Cocktail-exposed animals showed a significantly reduced ratio of new object contacts to total contacts (*p=*0.0279).

**Figure 6.**
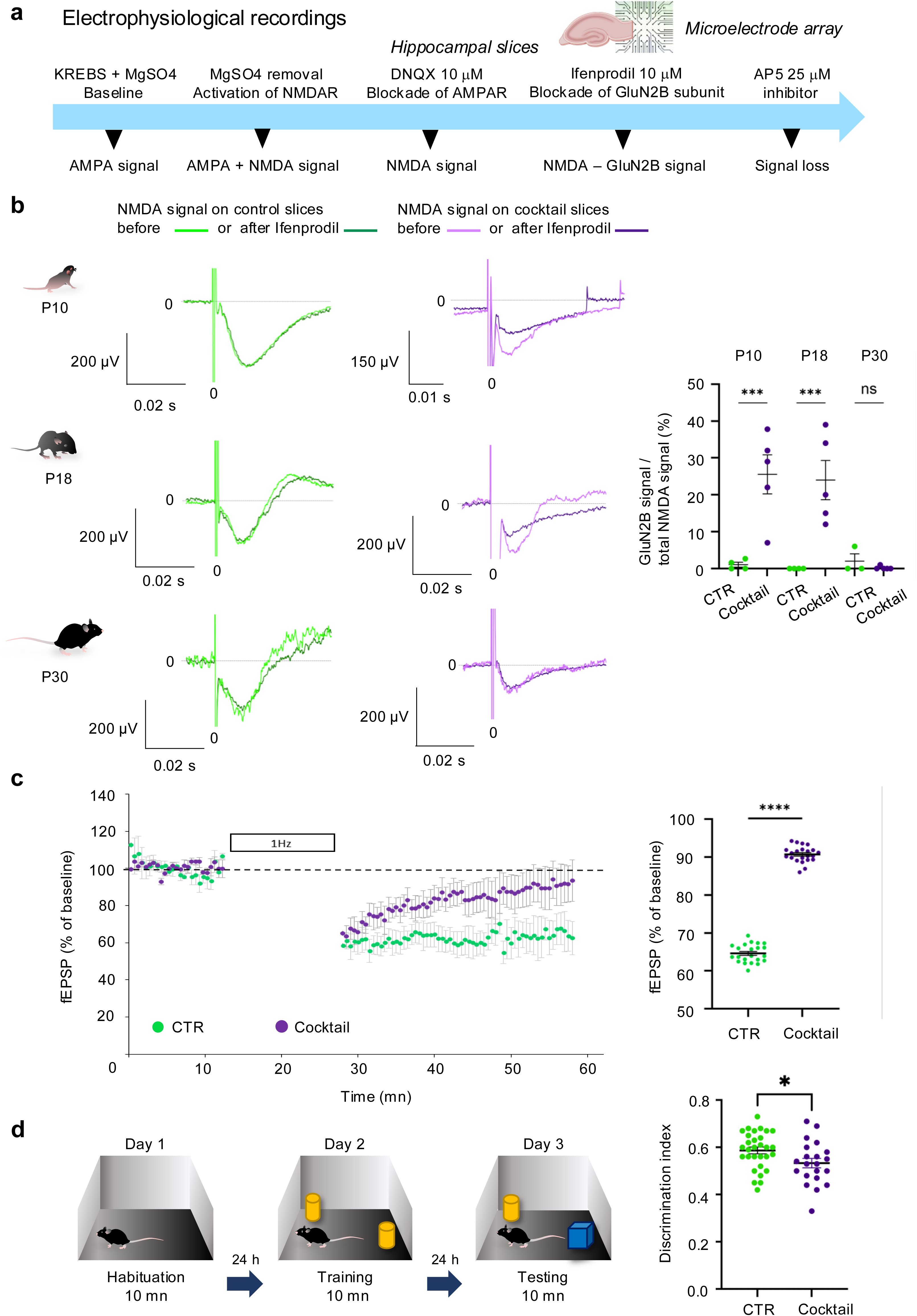
GluN2B-dependent signal is still present at P10 and P18 in acute hippocampal slices from cocktail-exposed animals and is associated with altered long-term depression (LTD). (**a**) Scheme of the electrophysiological protocol used for electrophysiological recordings on acute brain slices. Recordings of field excitatory post synaptic potentials (fEPSPs) were performed by transferring an hippocampal slice in a MEA continually perfused with extracellular solution to record AMPA receptor dependent fEPSPs. Magnesium ions-free extracellular solution was further applied in order to reveal NMDA receptor dependent synaptic signals. Then, DNQX (10 μM), an AMPA receptor antagonist was applied to the medium in order to isolate NMDA receptor dependent excitatory signals (fEPSP_NMDA_). Then, ifenprodil, a selective antagonist of the GluN2B subunit of the NMDA receptors was added to the medium to block the GluN2B signal. Finally, AP5 (25 μM), a non-selective NMDA receptor antagonist was applied to obtain complete inhibition of fEPSP_NMDA_. (**b**) Illustration of fEPSP_NMDA_ before (light green curve) and after (dark green curve) addition of Ifenprodil (10 μM) in slices from CTR mice. Representative fEPSP_NMDA_ before (light purple curve) and after (dark purple curve) addition of Ifenprodil (10 μM) in slices from cocktail-exposed mice. Each trace was obtained by averaging individual traces recorded over 2 minutes, before or after ifenprodil applicatio n. The decrease in signal corresponds to the blockade of the GluN2B receptor. The quantifica t io n of NMDA signal (%) due to the GluN2B subunit at P10, P18 and P30 is represented on the right graph. This percentage corresponds to the decrease in the total NMDA signal caused by the addition of ifenprodil. Results are presented as mean ± SEM. Statistical analysis: t-test after a Welch’s correction. (**c**) Mean curves of long-term depression (LTD) induced by a low-frequency stimulation protocol (1Hz for 15 minutes) in the hippocampus of 5 CTR mice (green dots) and 7 cocktail-exposed mice (purple dots). Quantification of LTD signal was performed by meaning LTD data per group (right graph: mean± SEM, t-test). (**d**) A novel object recognition test was performed in 3-month-old mice exposed or not to the cocktail of fungicides to evaluate their cognitive performance. On test session (D3), cocktail-exposed mice displayed reduced performance compared to CTR mice (mean± SEM, t-test). Asterisks: one symbol: p<0.05, two symbols: p<0.01; three symbols: p<0.001, four symbols: p<0.0001.

### 2.5 Alterations of developmental milestones in neonates exposed *in utero* to fungicide cocktail

We assessed the developmental consequences of early life exposure to fungicide cocktail by assessing multiple milestones in vehicle (CTR) and fungicide-exposed pups from P3 to P24 (Figure 7a). Body weight of cocktail-exposed pups was decreased compared to CTR animals from P8 to P24 (F_1,150_=8.3, p<0.01) (Figure 7b). As concerns sensory modalities, fungicide exposure delayed ear detachment at P3 (U=109.5, p<0.0001), but had no impact on eye opening and acoustic startle (Figure 7c). Regarding motricity, fungicide cocktail delayed the acquisit io n of righting reflex compared to controls (P4-P8; F_1,51_=5.0, p<0.05) without modifying ambulation (Figure 7d). In the open-field (P20), cocktail-exposed mice travelled a shorter distance (U=62.0, p<0.0001), possibly due to reduced activity, but entered more in the center of the arenas (U=113.5, p<0.01), suggesting low anxiety (Figure 7e). For the social repertoire, fungicide-exposed mice vocalized less at P3 but not at P7 (Figure 7f); at P10, cocktail-exposed pups showed no olfactory preference for maternal or foreign beddings compared to control animals which spend 80% of their time in the maternal litter (F_1,51_=34.6, p<0.0001) (Figure 7f). Finally, mice early exposed to fungicide cocktail displayed a severe deficit in social interactio n at P24, evidenced by strong decrease in the time spent in nose contact (CTR: 84s vs 17s in Cocktail; F_1,38_=650.3, p<0.0001), number of nose contact (CTR: 72 vs 33 Cocktail; F_1,38_=208.8, p<0.0001) and mean duration of a nose contact is divided by 2 (CTR: 1.16s vs 0.53s in Cocktail; F_1,38_=755.9, p<0.0001). Cocktail exposed mice made no paw contacts with their congener (CTR: 4.75 vs 0 in Cocktail; U=0.0, p<0.0001) and displayed a significant reduction in the number of following episodes (CTR: 10.95 vs 2.60 in Cocktail; U=1.5, p<0.0001). In contrast, these mice groomed more frequently (CTR: 2.35 vs 3.85 in Cocktail; F_1,38_=8.3, p<0.01), notably after social contact (CTR: 0.40 vs 3.45 in Cocktail; U=106.0, p<0.05), a sign of social discomfort. Consistent with decreased activity, they reared less during the test (CTR: 51.85 vs 27.70 in Cocktail; F_1,38_=85.2, p<0.0001). In conclusion, early life exposure to fungicide cocktail altered developmental trajectory in mice.

**Figure 7.**
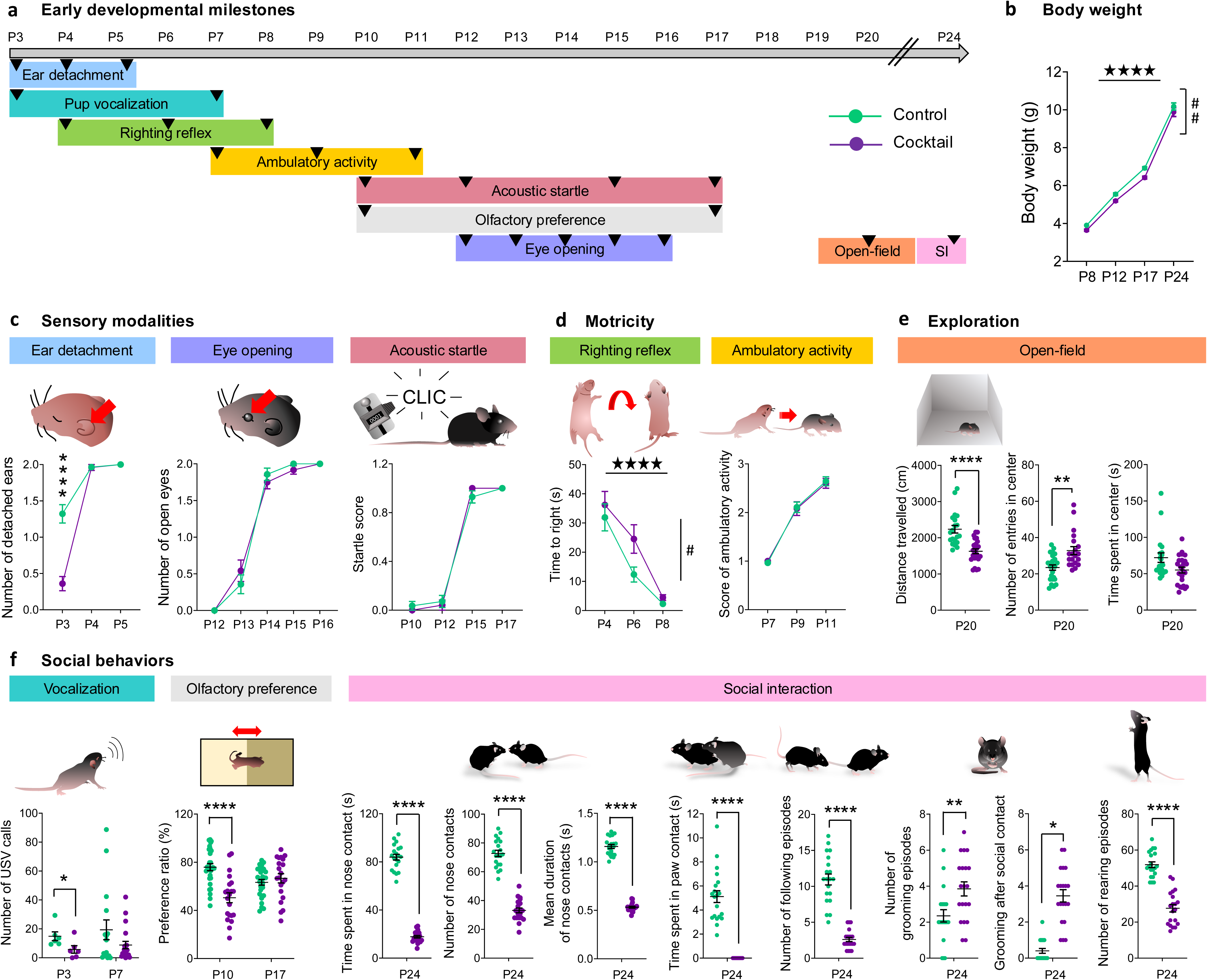
*In utero* exposure to a cocktail of fungicides impairs early developmental sensorial modalities, exploration ability and social behaviors. (**a**) Timeline of the early developmental tests performed in animals from P3 to P24, before weaning. Each colored label corresponds to a specific test and arrowheads indicate the timepoints when the tests were performed. All tests were performed successively in the same mice. (**b**) Cocktail-exposed mice showed reduced body weight compared to CTR mice between P8 and P24 (CTR n=28; Cocktail n=24, repeated measures ANOVA). (**c**) *Sensory modalities*: Fewer cocktail-exposed mice had their ears detached at P3 (CTR n=28; Cocktail n=24; Mann Whitney). (**d**) *Motricity:* righting reflex was delayed in cocktail-exposed mice (CTR n=28; Cocktail n=24; repeated measures ANOVA). (**e**) *Exploration*: mice exposed to the fungicide cocktail travelled a shorter distance in the open-field and entered more in the center of the apparatus than CTR mice (CTR n=20; Cocktail n=24; Mann-Whitney). (**f**) *Social behaviors:* cocktail-exposed mice emitted fewer ultrasonic calls than CTR at P3 (CTR n=6; Cocktail n=6; Mann-Whitney). At P10, cocktail-exposed mice failed to prefer maternal litter compared to the litter of an unfamiliar dam (CTR n=28; Cocktail n=24; unpaired t-test). At P24, mice exposed to fungicide cocktail showed a severe reduction in social interaction parameters associated with increased number of grooming episodes, notably after social contact (CTR n=20 with 10 males and 10 females; Cocktail n=20 with 10 males and 10 females; unpaired t-test or Mann-Whitney). Results are shown as scatter plots and/or mean ± sem. Asterisks: significant difference with CTR group (t-test or Mann Whitney); stars: time effect; hash: fungicide effect (repeated measures ANOVA); one symbol: p<0.05, two symbols: p<0.01; three symbols: p<0.001, four symbols: p<0.0001.

### 2.6 Young adult mice exposed *in utero* to fungicide cocktail exhibit autism-like phenotype

Fungicide-exposed and CTR mice underwent a series of behavioral tests at young adult stage (2-month-old) (Figure 8a). First, direct social interaction tests were done using pairs of animals of same sex, age and treatment, that have never met before (CTR n=16 with 8 males and 8 females; Cocktail n=16 with 8 males and 8 females) in an open field arena, during 10 min (Figure 8b). Regarding the social repertoire, the severe deficit in social interaction observed in cocktail-exposed mice at P24 was maintained in young adults, with decreased time spent in nose contact (CTR: 61s vs 11s in Cocktail; F_1,30_=619.7, p<0.0001), number (CTR: 52 vs 18 Cocktail; F_1,30_=276.6, p<0.0001) and mean duration of a nose contact (CTR: 1.18s vs 0.64s Cocktail; U=0.0, p<0.0001), absence of paw contact (CTR: 3.47 vs 0 in Cocktail; U=0.0, p<0.0001) and decreased number of following episodes (CTR: 5.87 vs 1.56 in Cocktail; F_1,30_=93.6, p<0.0001). The number of grooming episodes in cocktail-exposed mice was increased after a social contact only (CTR: 0.25 vs 1.87 in Cocktail; U=12.0, p<0.0001). Social preference and social novelty preference tests were performed in a 3-chamber apparatus, with 2 sessions of 10 min (Figure 8b). In the first phase of the three-chamber test, fungicide-exposed mice displayed a deficit in social preference. Indeed, they spent more time in nose contact (Treatment x Stimulus: F_1,30_=199.6, p<0.0001) and made more frequent (T x S: F_1,30_=48.3, p<0.0001) and longer (T x S: F_1,30_=1175.6, p<0.0001) contacts with the toy instead of the living mouse. The social preference ratio for the mouse is highly significant in CTR group, spending 80% of their time with congeners versus 30% for cocktail exposed mice (F_1,30_=654.9, p<0.0001). In the second phase, to determine social novelty preference, the toy is replaced by a new mouse (N) that has never been in contact with all mice used in this test. For CTR groups the social novelty preference is marked for the novel animal (N) whereas the social preference is marked for the familiar one (F) in cocktail groups: cocktail-exposed mice spent more time in nose contact (T x S: F_1,30_=313.8, p<0.0001) and made more frequent (T x S: F_1,30_=89.1, p<0.0001) and longer (T x S: F_1,30_=1382.1, p<0.0001) contacts with the familiar than the stranger mouse, leading to severely decreased social novelty preference ratio (30% for Cocktail vs 80% in CTR; F_1,30_=729.8, p<0.0001). As concerns nociception, the tail immersion test was performed at 3 temperatures: 48, 50 and 52°C (CTR n=16 with 8 males and 8 females; Cocktail n=16 with 8 males and 8 females) (Figure 8c). Early exposure to fungicide cocktail decreased the latency to flick at 48°C (F_1,30_=18.3, p<0.001) but increased this latency at 52°C (U=3.5, p<0.0001). For stereotypies/perseveration, fungicide exposure increased spontaneous digging (F_1,30_=4.2, p<0.05) and head shaking (U=32.5, p<0.001) in mice. Alternation in the different arms of the Y-maze test was scored to evaluate perseveration behavior by measuring spontaneous alternations (SPA), alternate arm returns (AAR) and same arm returns (SAR) (Figure 8d). Mice exposed to the cocktail of fungicides showed decreased spontaneous alternation (F_1,30_=144.7, p<0.0001) as a result of increased perseverative AAR (F_1,30_=38.5, p<0.0001) and SAR entries (U=1.0, p<0.0001), whereas the total number of entries in arms is unchanged. Finally, fungicide-exposed mice displayed exacerbated anxiety-like behavior in the novelty-suppressed feeding test, as shown by increased latency to eat (U=16.0, p<0.0001) (Figure 8e). Therefore, early-life exposure to the fungicide cocktail resulted in ASD-like social deficits and stereotyped behaviors in young mice, a phenotype that persisted into adulthood. At the adult stage, exposed mice also displayed altered nociceptive thresholds and increased anxiety, two frequent comorbidities of ASD^[38]^.

**Figure 8.**
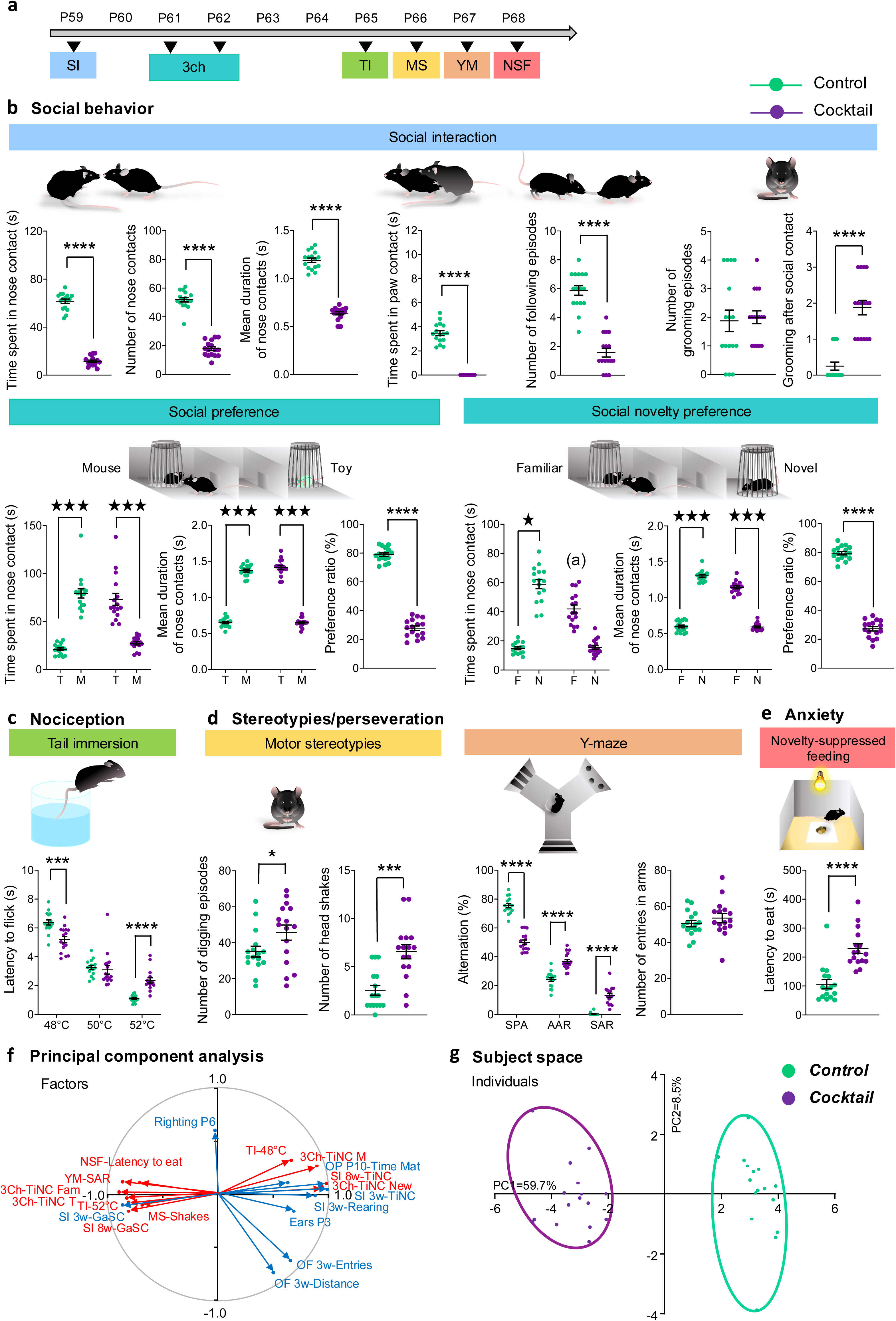
*In utero* exposure to a cocktail of fungicides impairs social interactions and increases perseverative behaviors in adult mice. (**a**) Timeline of behavioral tests performed in young adult mice from P59 to P68. Each colored label corresponds to a specific test. All tests were performed successively in the same mice (CTR n=16 with 8 males and 8 females; Cocktail n=16 with 8 males and 8 females). (**b**) *Social behavior*: Direct social interaction, social preference and social novelty preference were done at D59, D61 and D62, respectively. Social interaction tests were done using pairs of animals of same sex, age and treatment, that have never met before. The deficit in social interaction observed in cocktail-exposed mice was maintained at adult age (P59), with a general decrease in all parameters except for an increase in the number of grooming episodes after social contact (one-way ANOVA). Consistent with this, both social preference and social novelty preference (P61-62) were decreased in cocktail-exposed mice (repeated measures ANOVA). (**c**) *Nociception*: in the tail immersion test, mice exposed to the cocktail of fungicides (P65) displayed lower latency to flick at 48°C, but increased latency at 52°C (One-way ANOVA or Mann Whitney). (**d**) *Stereotypies/perseveration*: Cocktail-exposed mice displayed spontaneous stereotyped digging and head shaking behaviors. When exploring the Y-maze, they showed a decrease in spontaneous alternation (SPA), due to increased perseverative alternate arm returns (AAR) and same arm returns (SAR) (One-way ANOVA). (**e**) *Anxiety*: In the novelty-suppressed feeding test, cocktail-exposed mice took longer to eat in the center of the arena (Mann Whitney). Results are shown as scatter plots and/or mean ± sem. Asterisks: significant difference with CTR group (t-test); stars: time effect; hash: fungicide effect (repeated measures ANOVA); one asterisk: p<0.05, two asterisks: p<0.01; three asterisks: p<0.001, four asterisks: p<0.0001. (**f**) A PCA revealed that behavioral endpoints measured in pups (blue) and in adult mice (red) were highly correlated along PC1, driven by sociability parameters, while PC2 was mostly driven by activity (Factors). Individuals exposed to fungicides were clearly segregated from control mice in this analysis. 3Ch: three-chamber test; 3w: 3-week age; 8w: 8-week age; Ear: ear detachment; GaSC: grooming after social contact; MS: motor stereotypies; NSF: novelty-suppressed feeding test; OF: open-field; OP: olfactory preference; Righting: righting reflex in pups; SI: social interaction; TI: tail immersion; TiNC M/T: Time in nose contact with mouse/toy; TiNC Fam/New: Time in nose contact with familiar/new mouse; Time Mat: time in maternal bedding; YM-SAR: Y-maze test, same arm returns.

We used principal component analysis (PCA) to investigate whether behavioral deficits detected during early development in cocktail-exposed mice would correlate with ASD-like deficits in adulthood. In variable space (Figure 8f), most behavioral endpoints measured at 3-week age clustered with measures in adults along PC1 (59.7% of variance), driven by sociability parameters, with the exception of time to right (D6) and activity in the open-field, driven by locomotor activity along PC2 (8.5 % of variance). Thus, behavioral deficits observed in pups were highly correlated with altered responses observed later in young adults. In subject space (Figure 8g), PCA allowed to completely discriminate cocktail-exposed individuals from control mice along PC1, highlighting a clearly distinguishable phenotype between cocktail-exposed mice and controls.

### 2.7 Chronic exposure to fungicide cocktail residues induces changes in gene expression, including downregulation of GPCR purinergic receptors

Previously phenotyped mice were euthanized at adulthood (3 months of age) for gene expression analysis by qRT-PCR in three brain regions: the prefrontal cortex (PFC), nucleus accumbens (NAc) and ventral pallidum/olfactory tubercle (VP/Tu). We focused on genes coding for key players of glutamatergic neurotransmission and marker genes of striatal projection neurons (SPNs). Intensity map of differential gene expression (Figure 9a) show predominant over-expression in cocktail-exposed mice (cluster 1 (1)), except for a cluster of genes in the VP/Tu (cluster 2 (2)). When focusing on candidate genes, we unraveled upregulated expression of *Grin1* and *Grin2c* genes (encoding the GluN1 and GluN2C subunits of the NMDA receptor, respectively) in the NAc of fungicide-exposed mice. Interestingly, Grin1 and Grin2c displayed opposite regulation in the VP/Tu, with Grin1 upregulated and Grin2c downregulated. *Dlg3* (coding for PSD93) was upregulated in the PFC, whereas *Dlg4* (coding for PSD95) was upregulated in the NAc (Supplementary Figure S3). Among SPN marker genes, *Drd1* (dopamine D1 receptor) and *Adora2a* (adenosine 2A receptor) shared down-regulated expression in the VP/Tu (Figure 9a, Table S3).

**Figure 9.**
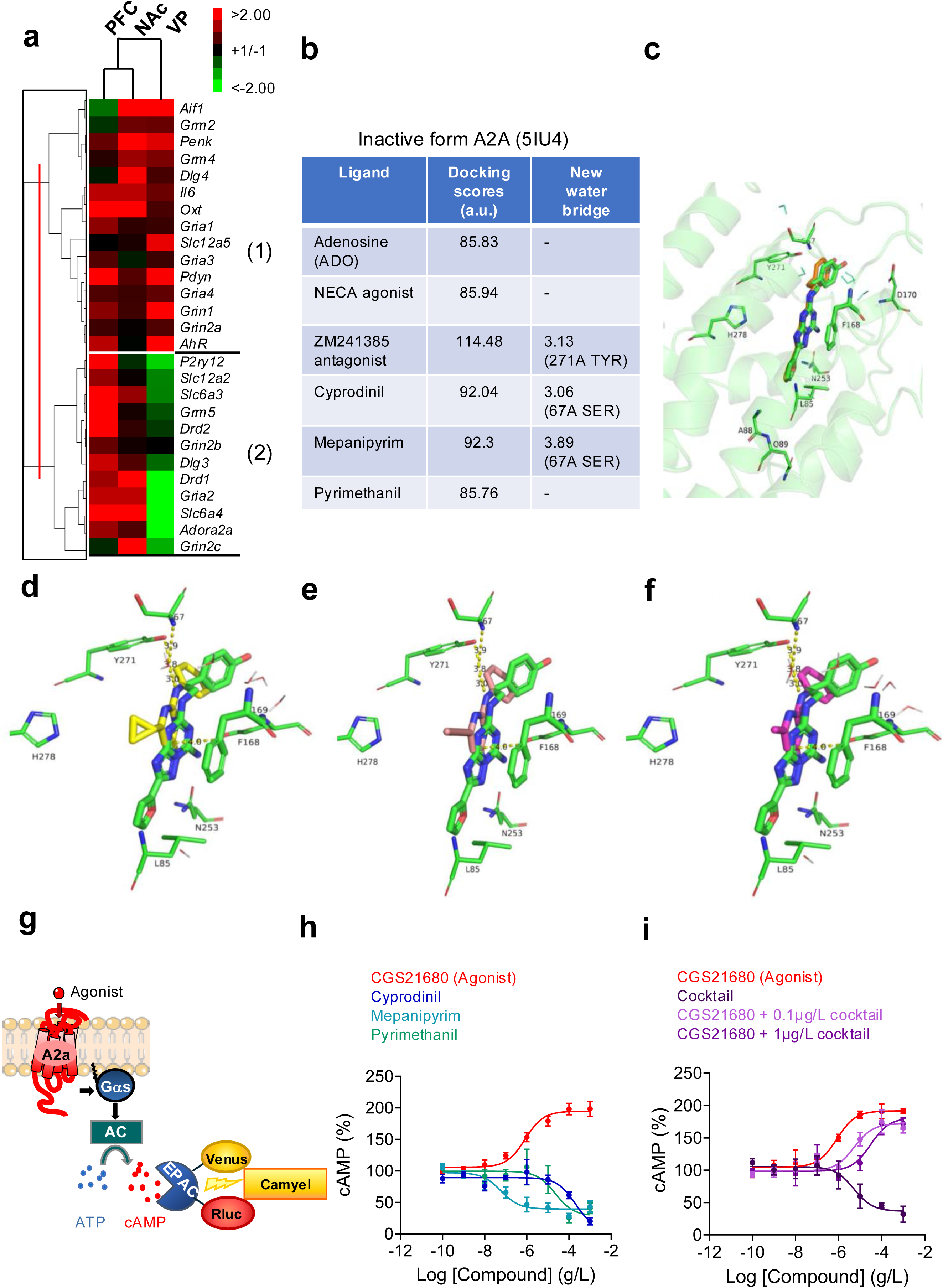
Anilinopyrimidine fungicides target the adenosine. A2A **receptor.** (**a**) The expression of 27 genes of interest coding for key players of glutamatergic neurotransmis s io n and marker genes of striatal projection neurons (SPNs) was assessed at 3-month age in the prefrontal cortex (PFC), nucleus accumbens (NAc) and ventral pallidum/olfactory tuber (VP/Tu) of previously phenotyped control and fungicide-exposed mice (CTR n = 8, Cocktail n = 8). In the heat map of gene expression per region, red color corresponds to upregulated expression and green color to down-regulated expression. Fungicide exposure globally increased gene expression, except for a cluster of genes under expressed in the VP/Tu, especially *Drd1, Gria2, Slc6a4* and *Adora2* coding for dopamine receptor, GluA2 subunit of AMPA receptor, serotonin transporter, and Adenosine A2A receptor. (**b**) Table of the docking scores, reflecting theoretical binding affinity, of the inactive form of A2A receptor (PDB 5IU4) with control ligands (adenosine: ADO; agonist: NECA; antagonist: ZM241385) and compared with anilinopyrimidine fungicides. Identification of novel water bridge involving residue SER 67 with cyprodinil and mepanipyrim. (**c**) Overview of the docking on A2A receptor (PDB 5IU4) and control with the antagonist ZM241385 form crystal (green sticks) and docking pose (orange sticks). Zoom-in views of docking poses show the ZM ligand (green) overlaid with (**d**) cyprodinil (yellow), (**e**) mepanipyrim (pale rose) and (**f**) pyrimethanil (pink sticks). Key residues of the A2A receptor are represented as thin green sticks, excepted F168, which is shown in thicker sticks. A new type of interaction is observed in the active site for cyprodinil and mepanipyrim, involving a water bridge between Y271 and S67, depicted as a yellow dotted line. (**g**) Principle of CAMYEL-based cAMP detection. A2A receptor activation stimula tes Gαs/AC-dependent cAMP production, monitored using the CAMYEL biosensor, a BRET-based EPAC probe composed of Renilla luciferase and Venus. cAMP binding induces a conformational change that alters the BRET signal. (**h**) CAMYEL concentration–respo nse curves following A2A receptor activation. Agonist CGS21680 increased cAMP signaling (EC₅₀ = 444 µg.L^-1^). Cyprodinil, mepanipyrim and pyrimethanil reduced the agonist-evoked CAMYEL response, with apparent IC₅₀ values of 201.4 μg.L⁻¹, 58.3 ng.L⁻¹ and 22.5 μg.L⁻¹, respectively. (**i**) CAMYEL assessment of a pesticide cocktail. Agonist CGS21680 increased cAMP signaling (EC₅₀ = 0.83 □M). The cocktail alone reduced basal CAMYEL signal (IC₅₀ = 51.7 μM) and shifted the CGS21680 dose–response curve rightward at 0.1 and 1 μg.L⁻¹, with apparent IC₅₀ values of 5.1 μM and 308.0 μM, respectively, indicating attenuation of agonist-induced signaling. Data are presented as mean ± S.E.M. from three independent experiments.

Based on transcriptomic data we hypothesized that one or more anilinopyrimidine compounds could interact with A2A receptor in adenosine ligand-binding pocket. We first compared their three-dimensional similarity to adenosine using the PubChem Score Matrix Service. Cyprodinil showed the highest similarity in molecular shape (Shape Tanimoto, ST = 81), followed by pyrimethanil (ST = 73) and mepanipyrim (ST = 72). In contrast, similarity in pharmacophoric features was low for all three compounds (Color Tanimoto, CT = 9). Accordingly, molecular docking studies were performed on the three fungicides using both inactive (Figure 9b-f) and active receptor^[39],[40]^ (Supplementary Figure S4) states, focusing on the adenosine binding site. Binding poses of the three anilinopyrimidines were fully superimposable in the orthosteric site of the inactive A2A receptor conformation. Their docking scores (reflecting their theoretical binding affinities) were comparable to or higher than that of the endogenous ligand adenosine (ADO) (86–92 vs 86 for ADO) (Figure 9b). Notably, for compounds with higher docking scores (cyprodinil and mepanipyrim), a distinct hydrogen bond involving a water molecule was observed with Ser67, differing from the water bridge formed between the antagonist ZM241385 and Tyr271. However, docking scores for the three anilinopyrimidines were lower in the active A2A receptor conformation than those of adenosine (ADO) and the agonist NECA (51–58 vs 65 for ADO) (Supplementary Figure S4). Accordingly, their binding poses were less well conserved and not fully superimposable in the active conformation.

Finally, we evaluated the effects of the individual fungicides on A2A receptor signalling in heterologous HEK293 cells using a state-of-the art BRET-based assay to monitor cAMP production following receptor activation by the selective A2A receptor agonist CGS21680^[41]^ (Figure 9g-h). Compounds concentrations were expressed as mass per volume to allow direct comparison with the concentrations used for the in vivo exposure paradigm and with drinking-water regulatory limits. Remarkably, all three fungicides reduced the constitutive cAMP signalling of the A2A receptor: cyprodinil (IC_50_ = 201.4 µg.L^-1^), mepanipyrim (IC_50_ = 58.3 ng.L^-1^) and pyrimethanil (IC_50_ = 22.5 µg.L^-1^). This reduction in basal receptor activity suggests that the fungicides behave as inverse agonists at the A2A receptor. The fungicide cocktail simila r ly inhibited basal A2A receptor signalling, with an IC₅₀ of 51.7 µg/L (Figure 9i), approximately 170-fold higher than the total concentration used for in vivo exposure (0.3 µg/L). Nevertheless, the cocktail altered agonist-induced signalling, producing a concentration-dependent rightward shift in the CGS21680 concentration-response curve. The apparent EC₅₀ increased from 446.8 µg/L to 2.73 mg/L at 0.3 µg/L cocktail (∼sixfold decrease in potency) and to 165.08 mg/L at 3 µg/L (Fig. 9i). Together, these results show that anilinopyrimidines alter *Adora2a* gene expression in vivo, inhibit constitutive A2A receptor activity and antagonize agonist-ind uced signalling.

## 3. Discussion

Epidemiological studies have shown that maternal exposure to pesticides through agricultural activities or residential proximity to crops is associated with an increased risk of neurodevelopmental disorders, including ASD and ADHD, in the offspring^[42],[43]^. The developing brain is particularly sensitive to environmental exposures, notably because of the incomplete maturation of the blood–brain barrier and immune system^[44]^. While repeated exposure to pollutants at high doses has been associated with adverse neurodevelopmental outcomes, the consequences of chronic exposure to doses considered safe for the general population remains poorly understood. Here, we show that developmental and life lo ng exposure to a mixture of anilinopyrimidine fungicides at a concentration below the regulatory limit for pesticide mixtures in drinking water (0.3 µg/L versus the 0.5 µg/L EU limit) disrupts cortical development, synaptic maturation and brain function. These alterations are associated with an atypical developmental trajectory and persistent deficits in social behavior, stereotyped behaviors, increased anxiety, altered nociception, and enhanced susceptibility to epileptifo rm activity, collectively supporting an ASD-like phenotype.

The cerebral cortex plays a central role in cognitive and emotional processing, and its development depends on tightly regulated neuronal migration to establish the characteristic six-layered cortical architecture. Genetic alterations or prenatal infections can disrupt this process, leading to cortical malformations, neurodevelopmental and neuropsychiatric disorders, and seizures^[45],[46]^. Our findings indicate that environmental pollutants may similarly interfere with cortical development. Using the multicolor IUE-based MAGIC Markers strategy, we identified several abnormalities specifically in fungicide-exposed animals, including deformation of CTIP2-positive deep cortical layers, clusters of disorganized cells and cobblestone-like structures containing neurons extending towards the pial surface. These abnormalities are consistent with impaired neuronal migration and resemble features described in developmental cortical disorders such as focal cortical dysplasia<u>^1^</u> and Walker–Warburg syndrome^[47]^. This interpretation is further supported by reduced levels of Doublecortin (DCX) at P10 in exposed animals. DCX plays a critical role in neuronal migration and cortical stratification, particula r ly during the multipolar-to-bipolar transition of migrating neurons^[48]^. Reduced DCX levels coincided with cortical layer deformation, cobblestone-like structures and the presence of abnormally multi-branched pyramidal neurons across cortical layers. Together, these findings indicate that developmental exposure to the fungicide mixture disrupts neuronal migration and cortical organization.

The structural abnormalities were accompanied by profound alterations in synaptic functio n. Malformations of cortical development are a major cause of epilepsy, and focal cortical dysplasia is frequently observed in children undergoing surgery for drug-resistant seizures^[49]^ . Both the cortex and hippocampus are also critically involved in neurodevelopmental disorders, epilepsy and ASD. We therefore investigated glutamatergic synaptic transmission in acute hippocampal slices from animals exposed to the fungicide cocktail in utero. Although the relative contribution of AMPA and NMDA receptors to basal synaptic transmission was not significantly altered, slices from exposed pups were substantially more prone to develop epileptiform field potentials, indicating increased network susceptibility to hyperexcitability. Notably, GluN2B-containing NMDA receptor-mediated synaptic responses persisted until at least P18 in exposed animals. During normal postnatal development, the progressive replacement of GluN2B-by GluN2A-containing NMDARs is a key step in glutamatergic synapse maturation and occurs during a critical period of synaptic development^[32],[33],[36]^. This developmental switch occurred as expected in control animals but was delayed in fungicide-exposed animals, in which GluN2B-mediated responses persisted beyond the normal developmental period. Such persistence may contribute to the increased susceptibility to epileptiform activity, given the proposed involvement of GluN2B-containing NMDARs in epileptogenesis associated with human cortical dysplasia^[30]^.

The delayed GluN2B-to-GluN2A switch was also associated with impaired synaptic plasticity. Reliable assessment of LTP was limited by the heterogeneity of responses and the frequent occurrence of epileptiform activity in slices from exposed animals. LTD could nevertheless be induced, but its maintenance was markedly impaired. LTD has an important role during early postnatal development, contributing to activity-dependent refinement of neuronal circuits, synaptic pruning, and processes underlying learning and memory^[33],[37]^. Alterations in the GluN2A/GluN2B balance have previously been associated with impaired hippocampal LTD and memory deficits. Consistent with these electrophysiological findings, exposed animals also performed less efficiently than controls in the novel object recognition test. The persistence of GluN2B-containing NMDARs beyond their normal developmental window, together with unstable LTD, is therefore consistent with delayed or impaired maturation of glutamatergic synapses following developmental exposure to the fungicide mixture.

These cellular and synaptic abnormalities provide a functional link between the early neurodevelopmental abnormalities and the subsequent behavioral phenotype. Fungicide-exposed pups displayed impaired ultrasonic communication, abnormal olfactory preference for the maternal litter, and pronounced deficits in social interaction. Social deficits persisted into late adolescence and adulthood, when exposed animals also showed impaired social and social-novelty preference, motor stereotypies and perseverative behaviors. At the adult stage, exposed mice additionally displayed increased anxiety and altered nociceptive thresholds, two frequent comorbidities of ASD in humans. Altogether, abnormalities in early developmental milesto nes, persistent social deficits and repetitive behaviors, together with anxiety, altered nociception and increased susceptibility to epileptiform activity, support an ASD-like phenotype following early exposure to the fungicide mixture^[50],[51]^.

This behavioral profile overlaps with phenotypes reported in rodent models exposed to other early environmental challenges, including maternal immune activation^[44]^ and prenatal exposure to valproic acid^[52]^ or herbicides^[12],[53]^. However, many previous toxicological studies have used exposure levels substantially higher than those typically encountered by the general population. For example, experimental studies examining the neurodevelopmental effects of glyphosate have used concentrations up to 980 mg/L in drinking water^[11]^ or oral gavage at 50 mg/kg body weight/day^[54]^, compared with an acceptable daily intake of 0.5 mg/kg body weight/day. By contrast, the anilinopyrimidine cocktail used here was administered at 0.3 µg/L, below the EU regulatory limit of 0.5 µg/L for pesticide mixtures in drinking water, corresponding to approximately 10 ng/kg body weight/day. These findings therefore raise questions about the potential neurodevelopmental consequences of chronic exposure to pesticide mixtures at concentrations currently considered safe for human consumption.

A key question is how exposure to anilinopyrimidine fungicides could produce such persistent developmental and synaptic alterations. Transcriptome analyses of young adult mice revealed changes in several genes involved in neuronal and synaptic function. In particular, *Adora2a*, *Drd1*, *Gria2* and *Slc6a4* were downregulated, across several brain regions, suggesting alterations in adenosinergic, dopaminergic, glutamatergic, and serotoninergic signaling. Reduced *Gria2* expression is particularly relevant to the electrophysiological phenotype, as GluA2-containing AMPA receptors are major determinants of excitatory synaptic transmis s io n and plasticity. In addition, *Dlg4*, which encodes the postsynaptic scaffolding protein PSD-95, was upregulated in the nucleus accumbens in exposed animals, consistent with the central role of PSD-95 in synaptic organization and neuropsychiatric disorders including ASD and schizophrenia^[55]^. In combination with the electrophysiological findings, these transcriptional changes point to a broader disruption of synaptic development and neurotransmission.

Among the genes modulated by fungicide exposure, *Adora2a* was of particular interest because it encodes the adenosine A2A receptor (A2AR), a G protein-coupled receptor with important functions in neurodevelopment^[56]^. Molecular modeling predicted direct interactions between A2AR and each of the three fungicides, cyprodinil, mepanipyrim, and pyrimethanil. These predictions were supported by in vitro pharmacological experiments showing that all three compounds behave as potent inverse agonists at A2AR. The convergence of transcripto mic, molecular-modeling, and pharmacological data therefore identifies A2AR signaling as a potential molecular target of anilinopyrimidines.

A2AR is particularly relevant in the context of developmental exposure. Although its expression becomes more restricted to specific neuronal populations in the adult brain, A2AR displays a broader distribution during embryonic and early postnatal development, including in the developing cortex and hippocampus. A2AR signaling has been implicated in neural progenitor proliferation, neuronal migration, axonal growth, synapse formation and activit y-dependent plasticity^[57],[56]^. This developmental expression pattern and diversity of functio ns suggest that A2AR may contribute to the establishment and maturation of neuronal networks before acquiring its more specialized neuro-modulatory functions in the mature brain. Our findings do not demonstrate that A2AR inhibition is sufficient to account for the neurodevelopmental phenotype. Instead, they suggest that altered A2AR signaling may contribute to at least two complementary aspects of the phenotype as illustrated in the graphical abstract:

One potential mechanism involves corticogenesis. A2AR signaling interacts functionally with developmental pathways that regulate neural progenitor development and neuronal migration^[56]^, including Wnt/β-catenin signaling. This pathway plays a major role in progenitor expansion, cell-fate determination and cortical lamination. Disruption of A2AR signaling during development could therefore contribute to the neuronal migration defects, abnormal cortical organization and reduced DCX levels observed in exposed animals. The molecular intermediates underlying this relationship remain to be established in our model, but these findings provide a plausible mechanistic link between the pharmacological targeting of A2AR and the early cortical abnormalities induced by fungicide exposure.

A second potential mechanism involves synaptic maturation and plasticity. Adenosine signaling through A2AR modulates synaptic transmission and activity-dependent plasticity and contributes to the capacity of neuronal networks to undergo and sustain long-term changes in synaptic strength^[57],[58]^. This provides a particularly relevant framework for our electrophysiological findings. The persistence of GluN2B-containing NMDARs beyond the normal developmental period, together with unstable LTD and increased susceptibility to epileptiform activity, points to delayed maturation and impaired plasticity of glutamatergic synapses in exposed animals. The concomitant downregulation of *Gria2*, coding for GluA2 subunit of AMPA receptor, further supports disruption of excitatory synaptic development. Whether altered A2AR signaling directly drives the delayed GluN2B-to-GluN2A switch or the LTD abnormalities remains to be determined.

These two mechanisms are not mutually exclusive. Impaired cortical development and delayed synaptic maturation could converge to disrupt the establishment, maturation and refinement of neuronal circuits, ultimately contributing to the persistent behavioral phenotype observed in exposed animals. This model is particularly relevant to ASD, in which alterations in cortical development, neuronal connectivity, and synaptic function are increasingly viewed as interconnected components of atypical developmental trajectories^[59]^. Consistent with a potential contribution of A2AR signaling, several reports showed that agonists of A2AR have beneficial effects on some core symptoms of ASD and their comorbidities, as epilepsy and anxiety^65^.

Overall, our findings support a model in which chronic exposure to environmentally relevant low doses of anilinopyrimidine fungicides alters neuronal migration and cortical organizat io n. These alterations are accompanied by delayed maturation of glutamatergic synapses, impaired synaptic plasticity and increased network excitability, ultimately associated with persistent ASD-like behavioral abnormalities. The convergence of transcriptomic, molecular-mode l ing and pharmacological evidence further identifies A2AR signaling as a plausible molecular interface linking anilinopyrimidine exposure to altered neurodevelopment.

## 5. Materials and Methods

### 5.1 Ethics statement

This project follows French national guidelines on animal experimentation and well-being and was approved by the French National Ethic Committee for Animal Experimenta t io n (APAFIS #19357-2019022020408583.v3 and APAFIS #38158-2022080316135358.v10). C57BL/6J WT mice were housed in a cage with a 12h light/12h dark cycle, water and food ad libitum (standard diet A04, SAFE Diets, France), at 23 ± 2°C and 53 ± 10% of relative humid ity. Igloos and laying were provided in each cage as enrichment. Animal studies are reported in compliance with the ARRIVE guidelines.

### 5.2 Animals’ treatment with fungicides

Fungicides used in this study were purchased from Sigma-Aldrich (France), cyprodinil (cat#34389), mepanipyrim (cat#33970) and pyrimethanil (cat#31577) (Figure 1a). Fungicides stock solutions were prepared as previously described^[20]^. Briefly, stock solutions were performed in glass vials (Wheaton sample vials, cat#Z256064, Sigma-Aldrich) at 1 mg/mL in pure DMSO (cat#D8418, Sigma-Aldrich). These solutions were diluted 1,000 times in distilled water to reach a final concentration of 1 mg/L. Solutions were then filtered (0.5 μm) and stored in glass bottles at 4°C until use. For animal treatment, stock solutions at 1 mg/L were further diluted 10,000 times in tap water to reach a final concentration of 0.1 μg/L. Animals were treated either with each fungicide alone at 0.1 μg/L, or with a cocktail of the three fungicides at 0.1 μg/L each. This concentration is at the limit or below the regulatory maximal limit dose allowed in drinking water (0.1 μg/L alone and 0.5 μg/L for the mixture)^[14]^. In parallel, a solution of pure DMSO (100%) was diluted 1,000 times in distilled water to a final concentration of 0.1%, filtered (0.5 μm) and stored in glass bottles at 4°C until use. To treat control vehicle animals, the stock solution of DMSO at 0.1% was further diluted 10,000 times in tap water (1x10^-5^ % corresponding to the equivalent volume of 1 fungicide). Thus, for an equivalent volume for 3 fungicides, the final percentage of DMSO used is 3x10^-5^%. Mouse water bottles were changed once a week for all groups and prepared from stock solutions as described above. One-month-old C57BL/6J WT female mice (G0) were treated with fungicides either alone or in cocktail, or with an equivalent volume of DMSO for 2 months (Figure 1a). After 2 months, groups of 2-3 females having received the same treatment were mated with males that had not previously been treated with fungicides. Female mice were still under fungicide treatment during gestation in order to mimic the vertical contamination with pesticide residues, and the treatment was continued until the offspring were sacrificed. The resulting progeny (G1) was sacrificed at different time points: 10, 18 and 30 days post-natal (P10, P18, P30) and their brains were collected to analyze distinct neurodevelopmental features. For corticogenesis studies (formation of cortical layers and organization of neurons and astrocytes), embryos were electroporated at E13.5 with Multi-Addressable Genome-Integrating Colors (MAGIC) markers^[21]^ and then post-natal day 10 (P10) newborns’ brain were analyzed (Figure 1b). To study the glutamatergic synaptic transmission, the Multi-Electrod Array (MEA) electrophysiological recording technique was used on brain slices at P10, P18 and P30 (Figure 1b). Several markers were studied through biochemical and histological analyses at P10 and P30, as well as gene expression analyses at 3 months old animals (Figure 1b). The number of mice used for each age is listed in the table (Figure 1c).

### 5.3 Tissue collection and preparation

Mice at different post-natal stages were sacrificed and brains were collected and weighted. For immunoblotting, the collected brains were immediately frozen in liquid nitrogen for biochemical analyses and stored at -80°C. Other brains were fixed in a commercial paraformaldehyde solution (Antigenfix, cat#P0014, Diapath) for 24 h (P10) and 48 h (P30) at 4°C. Brains were then rinsed in PBS, cryoprotected in a 30% sucrose solution for 4 days at 4°C and included in OCT (Tissue-Tek, cat#4583, Sakura Finetek). Brains embedded in OCT were quickly frozen in acetone chilled on dry ice and conserved at -80°C until use.

### 5.4 In utero electroporation and sample preparation

*In utero* electroporation (IUE) was performed as described previously in Loulier et al., (2020)^[21]^. Briefly, E13,5 pregnant females were anesthetized with Ketamine/Xylazine. A midline laparotomy was performed and uterine horns were exposed to allow embryo visualization. One microliter of DNA solution comprising plasmid expression vectors for the PB-CAG-Cytbow transgene, Cre recombinase and Piggybac transposase, mixed with sterile Fast Green dye (Sigma), was injected into the lateral ventricle of each embryo with a glass capillary. Three 50 ms pulses of 30 V were delivered to C57BL/6 embryos with 1.5–3 mm diameter Tweezertrodes positioned to target the dorsal wall of the lateral ventricle. Following surgery, the incision site was sutured (4-0, Ethicon) and pregnant females were allowed to wake up in a warming chamber. At 10 days post-natal, mice were weighed and perfused intracardia l ly with 4% paraformaldehyde. Dissected brains were post-fixed overnight (O/N) at 4°C in Antigenfix solution (Diapath), rinsed 3 times in 1X phosphate buffered saline (PBS) and embedded in 3% agarose. 200 µm-thick sections were cut with a vibratome (VT100S, Leica) in 1X PBS. Slices were subsequently processed for histological analysis: CTIP2 (1:500, Cat#ab18465, Abcam France) immunostaining; Hoechst 33342 (1: 2,000, cat#H3570, Invitrogen) staining. All samples were rinsed in 1X PBS, 3 times 5 min, and mounted using Vectashield medium (Cat#H-1900, Vector Labs) prior to imaging using confocal microscopy.

### 5.5 Imaging acquisition and quantification

Brain sections were imaged using a confocal microscope (Zeiss LSM880) and laser lines set up to separately excite mCerulean/mTurquoise2 (CFP), EYFP, tdTomato/mCherry (RFP) using 440, 514 and 561 nm, respectively. In addition, 633 nm and and 405 nm laser lines were used for AlexaFluor647 (CTIP2) and Hoechst staining, respectively. A 20x0,8 NA objective is used and XY field and Z-step size are adjusted according to the localization of electroporated cells and Nyquist criteria. The pinhole was set to 1 unit of Airy (AU) and the laser power adjusted to obtain optimal fluorescent signal-to-noise ratio without reaching saturation. 1,024x1,024 pixel Z-stacks were acquired using the motorized stage of the microscope, with a 10% overlap between adjacent stacks to subsequently enable mosaic reconstructions of the electroporated area or the region of interest. Neuron reconstruction was performed using Vaa3D software using neuron tracing plugin developed by Peng Lab ^[24]^.

### 5.6 Immunoblotting

The brains were weighed and resuspended (10 or 20% weight/volume) in lysis buffer (140 mM KCl, 10 mM NaH2PO4, 1.7 mM KH2PO4, 1 mM EDTA, H20 milliQ) containing 1X phosphatase inhibitor tablet (PhosSTOP, cat#04906845001, Roche) and 1X protease Inhibitor Cocktail (Mini cOmplete^TM^, cat#11836153001, Roche). Homogenization was carried out in microbead-containing tubes using a ribolyzer apparatus (FastPrep FP120, ThermoSavant). Brain homogenates, to which 2% SDS was added, were centrifuged at 14,000 rpm, 20 min at 4°C, to remove membrane debris. Protein concentration was measured on supernatants by BCA protein assay (Pierce), and samples were all normalized to the same concentration and diluted (1:1) in 2x loading buffer (4% SDS, 10% β-mercaptoethanol, 20% glycerol, 100 mM Tris-HCl pH 6.8, bromophenol blue). Samples were heated 10 min at 90°C before loading on a 4–15% gradient gels (Criterion^TM^ TGX^TM^ Precast gels, cat#5671085, Biorad). After protein transfer, PVDF membranes were blocked in 5% nonfat milk -TBS-Tween solution and were incubated overnight at 4°C with primary antibodies listed in Supplementary Table S1. Protein loading controls were performed with a mouse anti-β-actin. Membranes were revealed with a ChemiDoc™ MP Imager (Biorad) using the signal accumulation mode, and signals were quantified on the Image Lab software (version 6.0.1-Biorad). Briefly, signal was quantified by the Image Lab software and normalized by the quantification of β-actin signal.

### 5.7 Electrophysiology on brain slices

To evaluate neuronal network activities, the glutamatergic synaptic transmission was measured using Multi-Electrode Array (MEA) electrophysiological recordings performed on brain slices at P10, P18 and P30 (Figure 1b).

#### 5.7.1 Preparation of acute brain slices for recording

Mice were decapitated on ice for slices preparation. The whole brain was removed into ice-cold sucrose solution containing 2.5 mM KCl, 7mM MgCl_2_, 0.5 mM CaCl_2_, 1.2 mM NaH_2_PO_4_, 25mM NaHCO_3_, 11 mM Glucose, 234 mM sucrose. The whole brain was glued to a 3% agarose layer and oriented for coronal sections (250 μm) using a vibrating blade microtome (Leica VT1000S). The slices were then collected in the oxygenated (95%/5% O2/CO2) sucrose solution.

All slides containing hippocampal tissue were then placed in 1X Krebs solution (124 mM NaCl, 3 mM KCl, 26 mM NaHCO_3_, 1.25 mM NaH_2_PO_4_ (2H2O), 1 mM MgSO_4_, 1 mM D-Glucose, 2 mM CaCl_2_, 10 mM Hepes) and bubbled with 95%/5% O2/CO2 for at least 2h before use as described previously^[60]^. This solution was further used for electrophysiological recordings and referred as the ‘extracellular medium’.

#### 5.7.2 Electrophysiological recordings

Hippocampal slices were dissected and transferred to a Multi-Electrode Array (MEA-60; Multichannel systems) with 60 extracellular electrodes. The inter-electrode distance was 150 μm. A nylon mesh was positioned above the hippocampal slice to obtain a suitable electrical contact between the surface of the slice and the electrode array. Slices were continually perfused with the extracellular medium bubbled with O_2_/CO_2_ (95%/5%) and maintained at 32.5°C. Excitatory synaptic transmission was evoked in the dendritic subfield of CA1 hippocampal by afferent stimulation (Schaffer collaterals). Stimulation was achieved with an external stimulator by applying biphasic current pulses to one electrode of the array. Stimulation intensity (20 to 200 µA) and duration (70 to 100 µs) were adapted to avoid multiphasic responses that could appear upon a too-strong stimulation. Field excitatory postsynaptic potentials (fEPSPs) were recorded at the same time by the different electrodes of the array in the dendritic field of CA1 area. In order to analyze the contribution of postsynaptic NMDA receptors to synaptic transmission, magnesium ions-free extracellular medium was perfused to relieve NMDA receptor blockade. Then, AMPA /Kainate receptor antagonists (DNQX or GYKI 52466 both at 10 μM) were applied in order to isolate NMDA receptor-mediated responses (fEPSP_NMDA_). Further, in order to examine NMDA receptors subunit composition, selective antagonists were tested on fEPSP_NMDA_. More specifically, the presence of GluN2B subunit-containing NMDA receptors was assessed by applying ifenprodil (10 μM). Finally, AP5, a non-selective NMDA receptors antagonist (25 μM) was tested to obtain a complete blockade of NMDA receptor-mediated synaptic responses. Long term Depression (LTD) was triggered by applying a 1 Hertz stimulation to the slice for 15 minutes.

### 5.7.3 Analysis of electrophysiological recordings

Field EPSPs were recorded and analyzed with MC rack software (Multichannel systems). The ratio NMDA/AMPA was calculated by dividing the amplitude of fEPSP_NMDA_ by the amplitude of basal fEPSP. Further quantification of the areas of fEPSPs (area under curve; AUC) were performed with Matlab software. For each experiment, the contribution of GluN2B-containing NMDA postsynaptic receptors to excitatory transmission was thus calculated as follow: (AUC of fEPSP_NMDA_ – AUC of fEPSP_NMDA+ifenprodil_) / AUC of fESP_NMDA_. Each AUC represents the mean of 10 individuals AUC of fEPSPs recorded over 2 minutes either prior to (fEPSP_NMDA_) or during ifenprodil application (fEPSP_NMDA+ifenprodil_).

### 5.8 Behavioral studies

The acquisition of developmental milestones and reflexes ontology as well as early life behavior were assessed using procedures derived from Fox’s battery of tests and Wahlsten’s adaption of Fox’s tests^[61]^, Feather-Schussler tests^[62]^ and recently described in Vignon et al., (2025)^[63]^. Pups were tested individually, preferably in the morning (9 a.m.–1 p.m). To limit stress due to maternal separation, the time spent by the pup away from the mother and home cage was limited to the duration of each test. At the end of each test, the pup was immediately put back in its home cage. In young adults, social behaviors, nociception, stereotypies/perseveration, as well as anxiety tests have been performed as previously described^[38],[63],[50]^ .

### 5.9 Real-time quantitative PCR analysis

RNAs were extracted and purified using the Direct-Zol RNA MiniPrep kit (Zymo research, Irvine, USA). cDNA was synthetized using the ProtoScript II Reverse Transcriptase kit (New England BioLabs, Evry-Courcouronnes, France). qRT-PCR was performed in quadruplets on a CFX384 Touch Real-Time PCR Detection System (Biorad, Marnes-la-Coquette, France) using iQ-SYBR Green supermix (Bio-Rad) kit with 0.25 μl cDNA in a 12 μl final volume in Hard-Shell Thin-Wall 384-Well Skirted PCR Plates (Bio-rad). Gene-specific primers were designed using Primer3 software to obtain a 90- to 110-bp product; sequences are displayed in Table S2. Relative expression ratios were normalized to the level of actin and the 2^−ΔΔCt^ method was applied to evaluate differential expression level.

### 5.10 Functional studies with A2A receptor

CAMYEL assay was performed as previously described^[41]^. HEK293FT cells (Thermo Fisher Scientific, Waltham, MA, USA; RRID: CVCL_6911) were cultured in DMEM supplemented with 10% inactivated fetal bovine serum, 100 U ml⁻¹ penicillin, 0.1 mg ml⁻¹ streptomycin and 1 mM glutamine (Eurobio, France) at 37 °C in a humidified incubator containing 5% CO₂. Cells were routinely tested and confirmed negative for mycoplasma contamination. For functio nal assays, cells were seeded in 96-well plates and transiently transfected with 50 ng/well of human ADORA2A (hA2A) receptor cDNA (Missouri S and T cDNA resource center, USA) together with the CAMYEL biosensor using Metafectene PRO (Biontex, Germany), according to the manufacturer’s instructions. Assays were performed 36 h post-transfection, following 16 h serum-starvation. Then cells were stimulated with anilinopyrimidine compounds or reference agonists in PBS containing 250 μM IBMX (Tocris Bioscience, Bristol UK), 5 mM HEPES and 5 μM coelenterazine H. BRET signals were recorded for 35 min using a Mithras LB 943 microplate reader (Berthold Technologies). Mean BRET ratios obtained between 10 and 20 min after stimulation were used for analysis. Ligand responses were calculated relative to basal signal, and concentration–response curves were generated by nonlinear regression of normalized data.

### 5.11 Molecular docking of anilinopyrimidine compounds into A2A receptor

The binding capacities of anilinopyrimidine compounds toward the active and inactive conformations of the A2A receptor were predicted by molecular docking using the cryo-EM structures reported by Carpenter et al. (2016) (PDB: 5G53)^[39]^ and Segala et al. (2016)^[40]^ (PDB:5IU4), respectively. The predicted binding modes were then compared with those of the endogenous ligand adenosine (ADO), the agonist NECA, and the antagonist ZM241341. Docking was achieved using GOLD (Genetic Optimization for Ligand Docking, v2025.1, CCDC Software Limited) program^[64]^ by setting 50 runs of genetic algorithm for the search of binding poses and using the PLP (pairwise potentials) scoring function for ranking the best solutions. The targeting area was defined as a sphere of 10 Å radius around the natural ligand (ADO), NECA or ZM241341 and best-ranked binding poses were selected for detailed analysis of interactions between compounds and A2A residues using the PyMol Molecular Graphics System (v2.5, Schrödinger, LLC).

### 5.12 Statistical analyses

All data were initially analyzed on GraphPad Prism software (versions 8.0 and 9.0; GraphPad) using Shapiro-Wilk’s normality test. When data sets were normally distributed (p>0.05), statistical analyses were performed using parametric tests: multiple T-test (Holm-Sidak’s correction) or the One-way analysis of variance (ANOVA; Holm-Sidak’s post hoc analysis). However, when data sets were not normally distributed (p<0.05) statistical analyses were performed using a non-parametric test: Mann-Whitney U-test or Kruskal and Wallis.

A probability of 0.05 was defined as a significant difference for all statistical analyses. Welch’s correction was applied in cases where the variances differed significantly between groups. Groups are represented in a cloud of points with standard error mean.

*Principal component analysis.* A standard principal component analysis (PCA) was performed on behavioral data from early and adult behavioral phenotyping in mice exposed either to vehicle or fungicide cocktail, to reduce the dimensionality of the data set. We considered the two first extracted principal components (PC1 and PC2) as accounting for the most relevant variance in the data set and used them for schematic representation. Only behavioral parameters with coordinates >0.5 on at least one of these PC were kept for representation.

*qRT-PCR data*. As described previously^[65],[66]^, qRT-PCR data were transformed prior to statistical analysis to obtain a symmetrical distribution centred on 0, using the follow ing formula: if x<1, y=1-1/x; if x>1, y=x-1 (x: qPCR data; y: transformed data). Outliers over twice the standard deviation were excluded from calculations, as technical errors. Significance of qRT-PCR data was then assessed using a two-tailed t-test; an adjusted p-value was calculated using Benjamini-Hochberg correction for multiple testing. Unsupervised clustering analysis was performed on transformed qRT-PCR data using complete linkage with correlation distance (Pearson correlation) for drug, treatment and brain region (Cluster 3.0 and Treeview software).

### 5.13 AI-assisted technology in the writing process

During the preparation of this work the authors used ChatGPT 4o mini in order to correct the English of the manuscript. After using this tool, the authors reviewed and edited the content as needed and take full responsibility for the content of the publication.

## Abbreviations

Cypro: cyprodinil
Mepa: mepanipyrim
Pyri: pyrimethanil
EFSA: European Food Safety Agency
NMDA receptor: N-methyl-D-aspartic acid receptor
AMPA receptor: α-amino-3-hydroxy-5-methyl-4-isoxazolepropionic acid receptor
P3-P10-P30: post-natal day 3, 10 and 30.
GluN2B/2A: Glutamate receptor subunit e-2. N-methyl D-aspartate receptor subtype 2B/2A.
GluN1: Glutamate receptor ionotropic N-methyl D-aspartate receptor type1

## Conflict of interest

The authors declare that they have no conflict of interest.

## Acknowledgments

We acknowledge the imaging facility MRI, member of the national infrastructure France-BioImaging infrastructure supported by the French National Research Agency (ANR-10-INBS-04, «Investments for the future»). We thank Laetitia Hudecek, at the INM MRI Platform for her help in image acquisition. We acknowledge the RAM-Neuro animal facility, especially Denis Greuet and Antoine Picot for their assistance in animal behavior tests. Clemence Blanchot and Jason Durand for their help in USV recordings. We thank Nadine Mestre-Frances and Françoise Trousse for their histological advices, Anne Marcilhac for providing several antibodies for western blotting.

## Data availability

All data supporting the findings described in this manuscript are available in the article and in the Supplementary Information and from the corresponding author upon request. Source data are provided with this paper.

## Funding

This project was supported by several grants: The French National Research Agency (ANR) under the program “Investissement d’Avenir” ANR-16-IDEX-0006, I-Site MUSE STEMPest, ANR-22-CE34-0004 FunExpo, FRMENV202109013951, FRM EQU202103012576 Lehmann and the European Union (European project ExpoSignalz – GAP n°101156353). Views and opinions expressed are however those of the authors only, and do not necessary reflect those of the European. Neither the European Union, nor the granting authority can be held responsible for them. L.S.P. is the recipient of a fellowship ANR-16-IDEX-0006, I-Site MUSE STEMPest. A.V. is the recipient of a fellowship from Ecole Nationale Supérieure (ENS) Lyon. P.A.L. is the recipient of a fellowship from the French Ministry of Higher Education and Research. P.A.L., L.S.P., and A.V. have also received a 6 months fellowship from the Center of Excellence for Neurodegenerative diseases of Montpellier (CoEN) – CHU of Montpellier certified by AVIESAN as part of the PMND French plan (2014-2019).

## Authors’ contribution

V.P. conceived the study in its globality, participated in animal experiments, biochemica l studies, analyzed data and did interpretation. K.L. performed the *in utero* electroporatio n studies, participated in data analyses and interpretation. J.L.M and J.B. have designed and performed behavioral and transcriptomic studies, analyzed the data and did their interpretat io n. TL, JLM and JB performed pharmacology studies. L.S.P., A.V., U. A., P.A.L., D.T. and M.V. performed the experiments and analyzed data. LC performed molecular modeling studies of A2AR. V.P., JLM and L.S.P wrote the manuscript (first draft). All the authors read the manuscript, provided constructive criticisms and approved the final manuscript.

---

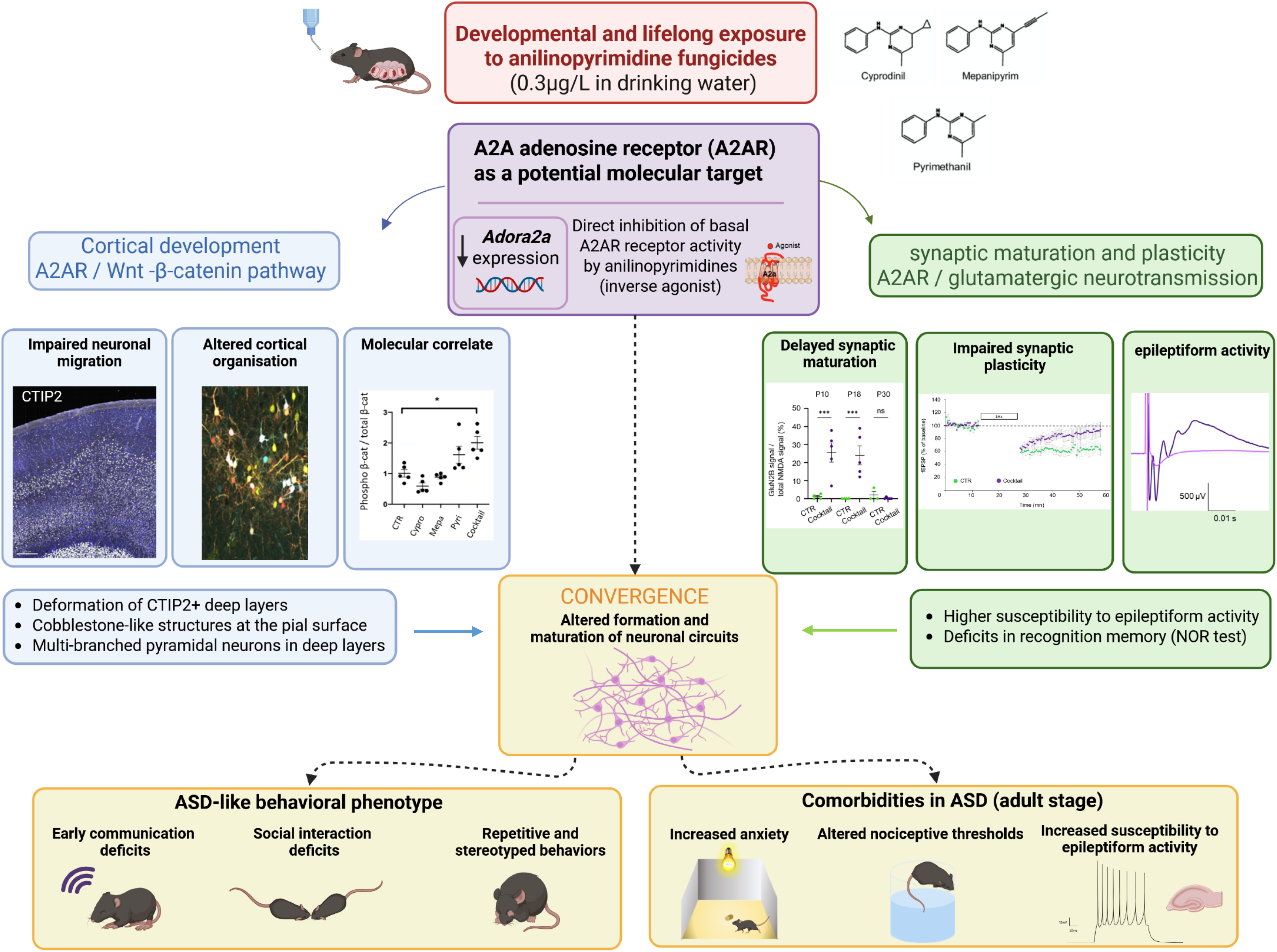

## Notes

### Competing Interest Statement

The authors have declared no competing interest.

## References

[1] . Diagnostic and statistical manual of mental disorders: DSM-5. 2013.

[2] Al-Beltagi, M., Autism medical comorbidities.World Journal of Clinical Pediatrics. 10. (3). 15–28. 2021. 10.5409/wjcp.v10.i3.15.

[3] Hallmayer, J. Genetic Heritability and Shared Environmental Factors Among Twin Pairs With Autism.Archives of General Psychiatry. 68. (11). 1095. 2011. 10.1001/archgenpsychiatry.2011.76.

[4] Chaste, P., Leboyer, M. Autism risk factors: genes, environment, and gene-environment interactions.Dialogues in Clinical Neuroscience. 14. (3). 281–292. 2012.

[5] Chen, J. A., Peñagarikano, O., Belgard, T. G., Swarup, V., Geschwind, D. H. The Emerging Picture of Autism Spectrum Disorder: Genetics and Pathology.Annual Review of Pathology: Mechanisms of Disease. 10. (1). 111–144. 2015. 10.1146/annurev-pathol-012414-040405.

[6] Rauh, V., Arunajadai, S., Horton, M., Perera, F., Hoepner, L., Barr, D. B., Whyatt, R. Seven-year neurodevelopmental scores and prenatal exposure to chlorpyrifos, a common agricultural pesticide.Environmental Health Perspectives. 119. (8). 1196–1201. 2011. 10.1289/ehp.1003160.

[7] Juntarawijit, Y., Chaichanawirote, U., Rakmeesri, P., Chairattanasakda, P., Pumyim, V., Juntarawijit, C. Chlorpyrifos and other pesticide exposure and suspected developmental delay in children aged under 5 years: a case-control study in Phitsanulok, Thailand. F1000Research. 9. 1501. 2020. 10.12688/f1000research.27874.5.

[8] Brown, A. S., Cheslack-Postava, K., Rantakokko, P., Kiviranta, H., Hinkka-Yli-Salomäki, S., McKeague, I. W., Surcel, H.-M., Sourander, A. Association of Maternal Insecticide Levels With Autism in Offspring From a National Birth Cohort.The American Journal of Psychiatry. 175. (11). 1094–1101. 2018. 10.1176/appi.ajp.2018.17101129.

[9] Ehrenstein, O. S. von, Ling, C., Cui, X., Cockburn, M., Park, A. S., Yu, F., Wu, J., Ritz, B. Prenatal and infant exposure to ambient pesticides and autism spectrum disorder in children: population based case-control study. BMJ (Clinical research ed.). 364. l962. 2019. 10.1136/bmj.l962.

[10] Samsel, A., Seneff, S. Glyphosate, pathways to modern diseases III: Manganese, neurological diseases, and associated pathologies.Surgical Neurology International. 6. 45. 2015. 10.4103/2152-7806.153876.

[11] Pu, Y., Yang, J., Chang, L., Qu, Y., Wang, S., Zhang, K., Xiong, Z., Zhang, J., Tan, Y., Wang, X., Fujita, Y., Ishima, T., Wan, D., Hwang, S. H., Hammock, B. D., Hashimoto, K. Maternal glyphosate exposure causes autism-like behaviors in offspring through increased expression of soluble epoxide hydrolase.Proceedings of the National Academy of Sciences. 117. (21). 11753–11759. 2020. 10.1073/pnas.1922287117.

[12] Laugeray, A., Herzine, A., Perche, O., HÃ©bert, B., Aguillon-Naury, M., Richard, O., Menuet, A., Mazaud-Guittot, S., LesnÃ©, L., Briault, S., Jegou, B., Pichon, J., MontÃ©cot-Dubourg, C., Mortaud, S. Pre- and Postnatal Exposure to Low Dose Glufosinate Ammonium Induces Autism-Like Phenotypes in Mice.Frontiers in Behavioral Neuroscience. 8. 2014. 10.3389/fnbeh.2014.00390.

[13] AirParif. Surveillance et information sur la qualité de l’air en Ile-de-France. 2016.

[14] Council of the European Union. Council directive 98/83EC of 3 November 1998 on the quality of water for human consumption.Official Journal of the European Communities. L.330. 32–54. 1998.

[15] ANSES. Evaluation des risques liés aux résidus de pesticides dans l’eau de distribution. Contribution à l’exposition alimentaire totale. 2013.

[16] EFSA. The 2017 European Union report on pesticide residues in food.17(6):5743. 2019.

[17] Nougadère, A., Sirot, V., Cravedi, J.-P., Vasseur, P., Feidt, C., Fussell, R. J., Hu, R., Leblanc, J.-C., Jean, J., Rivière, G., Sarda, X., Merlo, M., Hulin, M. Dietary exposure to pesticide residues and associated health risks in infants and young children - Results of the French infant total diet study.Environment International. 137. 105529. 2020. 10.1016/j.envint.2020.105529.

[18] Béranger, R., Hardy, E. M., Dexet, C., Guldner, L., Zaros, C., Nougadère, A., Metten, M.-A., Chevrier, C., Appenzeller, B. M. R. Multiple pesticide analysis in hair samples of pregnant French women: Results from the ELFE national birth cohort.Environment International. 120. 43–53. 2018. 10.1016/j.envint.2018.07.023.

[19] Corrales Vargas, A., Peñaloza Castañeda, J., Rietz Liljedahl, E., Mora, A. M., Menezes-Filho, J. A., Smith, D. R., Mergler, D., Reich, B., Giffin, A., Hoppin, J. A., Lindh, C. H., Wendel de Joode, B. van. Exposure to common-use pesticides, manganese, lead, and thyroid function among pregnant women from the Infants’ Environmental Health (ISA) study, Costa Rica. The Science of the Total Environment. 810. 151288. 2022. 10.1016/j.scitotenv.2021.151288.

[20] Wang, Y., Lafon, P.-A., Salvador-Prince, L., Gines, A. R., Trousse, F., Torrent, J., Prevostel, C., Crozet, C., Liu, J., Perrier, V. Prenatal exposure to low doses of fungicides corrupts neurogenesis in neonates.Environmental Research. 195. 110829. 2021. 10.1016/j.envres.2021.110829.

[21] Loulier, K., Barry, R., Mahou, P., Le Franc, Y., Supatto, W., Matho, K. S., Ieng, S., Fouquet, S., Dupin, E., Benosman, R., Chédotal, A., Beaurepaire, E., Morin, X., Livet, J. Multiplex Cell and Lineage Tracking with Combinatorial Labels.Neuron. 81. (3). 505–520. 2014. 10.1016/j.neuron.2013.12.016.

[22] Clavreul, S., Abdeladim, L., Hernández-Garzón, E., Niculescu, D., Durand, J., Ieng, S.-H., Barry, R., Bonvento, G., Beaurepaire, E., Livet, J., Loulier, K. Cortical astrocytes develop in a plastic manner at both clonal and cellular levels.Nature Communications. 10. (1). 4884. 2019. 10.1038/s41467-019-12791-5.

[23] Hevner, R. F. Layer-Specific Markers as Probes for Neuron Type Identity in Human Neocortex and Malformations of Cortical Development:Journal of Neuropathology and Experimental Neurology. 66. (2). 101–109. 2007. 10.1097/nen.0b013e3180301c06.

[24] Peng, H., Bria, A., Zhou, Z., Iannello, G., Long, F. Extensible visualization and analysis for multidimensional images using Vaa3D.Nature Protocols. 9. (1). 193–208. 2014. 10.1038/nprot.2014.011.

[25] Adachi, K., Mirzadeh, Z., Sakaguchi, M., Yamashita, T., Nikolcheva, T., Gotoh, Y., Peltz, G., Gong, L., Kawase, T., Alvarez-Buylla, A., Okano, H., Sawamoto, K. Beta-catenin signaling promotes proliferation of progenitor cells in the adult mouse subventricular zone.Stem Cells (Dayton, Ohio). 25. (11). 2827–2836. 2007. 10.1634/stemcells.2007-0177.

[26] Adachi, K., Mirzadeh, Z., Sakaguchi, M., Yamashita, T., Nikolcheva, T., Gotoh, Y., Peltz, G., Gong, L., Kawase, T., Alvarez-Buylla, A., Okano, H., Sawamoto, K. Beta-catenin signaling promotes proliferation of progenitor cells in the adult mouse subventricular zone.Stem Cells (Dayton, Ohio). 25. (11). 2827–2836. 2007. 10.1634/stemcells.2007-0177.

[27] Yu, X., Malenka, R. C. Beta-catenin is critical for dendritic morphogenesis.Nature Neuroscience. 6. (11). 1169–1177. 2003. 10.1038/nn1132.

[28] Ramos-Fernández, E., Tapia-Rojas, C., Ramírez, V. T., Inestrosa, N. C. Wnt-7a Stimulates Dendritic Spine Morphogenesis and PSD-95 Expression Through Canonical Signaling.Molecular Neurobiology. 56. (3). 1870–1882. 2019. 10.1007/s12035-018-1162-1.

[29] Hansen, K. B., Wollmuth, L. P., Bowie, D., Furukawa, H., Menniti, F. S., Sobolevsky, A. I., Swanson, G. T., Swanger, S. A., Greger, I. H., Nakagawa, T., McBain, C. J., Jayaraman, V., Low, C.-M., Dell’Acqua, M. L., Diamond, J. S., Camp, C. R., Perszyk, R. E., Yuan, H., Traynelis, S. F. Structure, Function, and Pharmacology of Glutamate Receptor Ion Channels.Pharmacological Reviews. 73. (4). 1469–1658. 2021. 10.1124/pharmrev.120.000131.

[30] Möddel, G., Jacobson, B., Ying, Z., Janigro, D., Bingaman, W., González-Martínez, J., Kellinghaus, C., Prayson, R. A., Najm, I. M. The NMDA receptor NR2B subunit contributes to epileptogenesis in human cortical dysplasia.Brain Research. 1046. (1-2). 10–23. 2005. 10.1016/j.brainres.2005.03.042.

[31] Monyer, H., Burnashev, N., Laurie, D. J., Sakmann, B., Seeburg, P. H. Developmental and regional expression in the rat brain and functional properties of four NMDA receptors.Neuron. 12. (3). 529–540. 1994. 10.1016/0896-6273(94)90210-0.

[32] Liu, X.-B., Murray, K. D., Jones, E. G. Switching of NMDA receptor 2A and 2B subunits at thalamic and cortical synapses during early postnatal development.The Journal of Neuroscience: The Official Journal of the Society for Neuroscience. 24. (40). 8885–8895. 2004. 10.1523/JNEUROSCI.2476-04.2004.

[33] Yashiro, K., Philpot, B. D. Regulation of NMDA receptor subunit expression and its implications for LTD, LTP, and metaplasticity.Neuropharmacology. 55. (7). 1081–1094. 2008. 10.1016/j.neuropharm.2008.07.046.

[34] Bartlett, T. E., Bannister, N. J., Collett, V. J., Dargan, S. L., Massey, P. V., Bortolotto, Z. A., Fitzjohn, S. M., Bashir, Z. I., Collingridge, G. L., Lodge, D. Differential roles of NR2A and NR2B-containing NMDA receptors in LTP and LTD in the CA1 region of two-week old rat hippocampus.Neuropharmacology. 52. (1). 60–70. 2007. 10.1016/j.neuropharm.2006.07.013.

[35] Kellermayer, B., Ferreira, J. S., Dupuis, J., Levet, F., Grillo-Bosch, D., Bard, L., Linarès-Loyez, J., Bouchet, D., Choquet, D., Rusakov, D. A., Bon, P., Sibarita, J.-B., Cognet, L., Sainlos, M., Carvalho, A. L., Groc, L. Differential Nanoscale Topography and Functional Role of GluN2-NMDA Receptor Subtypes at Glutamatergic Synapses.Neuron. 100. (1). 106–119.e7. 2018. 10.1016/j.neuron.2018.09.012.

[36] Shipton, O. A., Paulsen, O. GluN2A and GluN2B subunit-containing NMDA receptors in hippocampal plasticity.Philosophical Transactions of the Royal Society of London. Series B, Biological Sciences. 369. (1633). 20130163. 2014. 10.1098/rstb.2013.0163.

[37] Drissi, I., Deschamps, C., Fouquet, G., Alary, R., Peineau, S., Gosset, P., Sueur, H., Marcq, I., Debuysscher, V., Naassila, M., Vilpoux, C., Pierrefiche, O. Memory and plasticity impairment after binge drinking in adolescent rat hippocampus: GluN2A/GluN2B NMDA receptor subunits imbalance through HDAC2.Addiction Biology. 25. (3). e12760. 2020. 10.1111/adb.12760.

[38] Spencer, C. M., Alekseyenko, O., Serysheva, E., Yuva-Paylor, L. A., Paylor, R. Altered anxiety-related and social behaviors in the Fmr1 knockout mouse model of fragile X syndrome.Genes, Brain, and Behavior. 4. (7). 420–430. 2005. 10.1111/j.1601-183X.2005.00123.x.

[39] Carpenter, B., Nehmé, R., Warne, T., Leslie, A. G. W., Tate, C. G. Structure of the adenosine A2A receptor bound to an engineered G protein.Nature. 536. (7614). 104–107. 2016. 10.1038/nature18966.

[40] Segala, G., Bennesch, M. A., Pandey, D. P., Hulo, N., Picard, D. Monoubiquitination of Histone H2B Blocks Eviction of Histone Variant H2A.Z from Inducible Enhancers.Molecular Cell. 64. (2). 334–346. 2016. 10.1016/j.molcel.2016.08.034.

[41] Laboute, T., Gandía, J., Pellissier, L. P., Corde, Y., Rebeillard, F., Gallo, M., Gauthier, C., Léauté, A., Diaz, J., Poupon, A., Kieffer, B. L., Le Merrer, J., Becker, J. A. The orphan receptor GPR88 blunts the signaling of opioid receptors and multiple striatal GPCRs.eLife. 9. e50519. 2020. 10.7554/eLife.50519.

[42] Ongono, J. S., Béranger, R., Baghdadli, A., Mortamais, M. Pesticides used in Europe and autism spectrum disorder risk: can novel exposure hypotheses be formulated beyond organophosphates, organochlorines, pyrethroids and carbamates? - A systematic review.Environmental Research. 187. 109646. 2020. 10.1016/j.envres.2020.109646.

[43] Shelton, J. F., Geraghty, E. M., Tancredi, D. J., Delwiche, L. D., Schmidt, R. J., Ritz, B., Hansen, R. L., Hertz-Picciotto, I. Neurodevelopmental Disorders and Prenatal Residential Proximity to Agricultural Pesticides: The CHARGE Study.Environmental Health Perspectives. 122. (10). 1103–1109. 2014. 10.1289/ehp.1307044.

[44] Tartaglione, A. M., Villani, A., Ajmone-Cat, M. A., Minghetti, L., Ricceri, L., Pazienza, V., De Simone, R., Calamandrei, G. Maternal immune activation induces autism-like changes in behavior, neuroinflammatory profile and gut microbiota in mouse offspring of both sexes.Translational Psychiatry. 12. (1). 384. 2022. 10.1038/s41398-022-02149-9.

[45] Buchsbaum, I. Y., Cappello, S. Neuronal migration in the CNS during development and disease: insights from in vivo and in vitro models.Development (Cambridge, England). 146. (1). dev163766. 2019. 10.1242/dev.163766.

[46] Romero, D., N, B.-B., F, F. Genetics and mechanisms leading to human cortical malformations.Seminars in cell & developmental biology. 76. 2018. 10.1016/j.semcdb.2017.09.031.

[47] Judas, M., Sedmak, G., Rados, M., Sarnavka, V., Fumić, K., Willer, T., Gross, C., Hehr, U., Strahl, S., Cuk, M., Barić, I. POMT1-associated walker-warburg syndrome: a disorder of dendritic development of neocortical neurons.Neuropediatrics. 40. (1). 6–14. 2009. 10.1055/s-0029-1224099.

[48] Volvert, M.-L. MicroRNA Targeting of CoREST Controls Polarization of Migrating Cortical Neurons.16.

[49] Blumcke, I., Spreafico, R., Haaker, G., Coras, R., Kobow, K., Bien, C. G., Pfäfflin, M., Elger, C., Widman, G., Schramm, J., Becker, A., Braun, K. P., Leijten, F., Baayen, J. C., Aronica, E., Chassoux, F., Hamer, H., Stefan, H., Rössler, K., Thom, M., Walker, M. C., Sisodiya, S. M., Duncan, J. S., McEvoy, A. W., Pieper, T., Holthausen, H., Kudernatsch, M., Meencke, H. J., Kahane, P., Schulze-Bonhage, A., Zentner, J., Heiland, D. H., Urbach, H., Steinhoff, B. J., Bast, T., Tassi, L., Lo Russo, G., Özkara, C., Oz, B., Krsek, P., Vogelgesang, S., Runge, U., Lerche, H., Weber, Y., Honavar, M., Pimentel, J., Arzimanoglou, A., Ulate-Campos, A., Noachtar, S., Hartl, E., Schijns, O., Guerrini, R., Barba, C., Jacques, T. S., Cross, J. H., Feucht, M., Mühlebner, A., Grunwald, T., Trinka, E., Winkler, P. A., Gil-Nagel, A., Toledano Delgado, R., Mayer, T., Lutz, M., Zountsas, B., Garganis, K., Rosenow, F., Hermsen, A., Oertzen, T. J. von, Diepgen, T. L., Avanzini, G. Histopathological Findings in Brain Tissue Obtained during Epilepsy Surgery.New England Journal of Medicine. 377. (17). 1648–1656. 2017. 10.1056/NEJMoa1703784.

[50] Silverman, J. L., Yang, M., Lord, C., Crawley, J. N. Behavioural phenotyping assays for mouse models of autism.Nature Reviews. Neuroscience. 11. (7). 490–502. 2010. 10.1038/nrn2851.

[51] Argyropoulos, A., Gilby, K. L., Hill-Yardin, E. L. Studying Autism in Rodent Models: Reconciling Endophenotypes with Comorbidities.Frontiers in Human Neuroscience. 7. 2013. 10.3389/fnhum.2013.00417.

[52] Hughes, E. M., Calcagno, P., Clarke, M., Sanchez, C., Smith, K., Kelly, J. P., Finn, D. P., Roche, M. Prenatal exposure to valproic acid reduces social responses and alters mRNA levels of opioid receptor and pre-pro-peptide in discrete brain regions of adolescent and adult male rats.Brain Research. 1732. 146675. 2020. 10.1016/j.brainres.2020.146675.

[53] Ait-Bali, Y., Ba-M’hamed, S., Gambarotta, G., Sassoè-Pognetto, M., Giustetto, M., Bennis, M. Pre- and postnatal exposure to glyphosate-based herbicide causes behavioral and cognitive impairments in adult mice: evidence of cortical ad hippocampal dysfunction.Archives of Toxicology. 94. (5). 1703–1723. 2020. 10.1007/s00204-020-02677-7.

[54] Oliveira, M. A. L. de, Rojas, V. C. T., Sá, J. C. de, Novais, C. O. de, Silva, M. S., Almeida Paula, H. A. de, Kirsten, T. B., Bernardi, M. M., Pinheiro, L. C., Giusti-Paiva, A., Vilela, F. C. Perinatal exposure to glyphosate-based herbicides induced neurodevelopmental behaviors impairments and increased oxidative stress in the prefrontal cortex and hippocampus in offspring.International Journal of Developmental Neuroscience: The Official Journal of the International Society for Developmental Neuroscience. 82. (6). 528–538. 2022. 10.1002/jdn.10207.

[55] Hormozdiari, F., Penn, O., Borenstein, E., Eichler, E. E. The discovery of integrated gene networks for autism and related disorders.Genome Research. 25. (1). 142–154. 2015. 10.1101/gr.178855.114.

[56] Rimbert, S., Moreira, J. B., Xapelli, S., Lévi, S. Role of purines in brain development, from neuronal proliferation to synaptic refinement.Neuropharmacology. 237. 109640. 2023. 10.1016/j.neuropharm.2023.109640.

[57] Alçada-Morais, S., Gonçalves, N., Moreno-Juan, V., Andres, B., Ferreira, S., Marques, J. M., Magalhães, J., Rocha, J. M. M., Xu, X., Partidário, M., Cunha, R. A., López-Bendito, G., Rodrigues, R. J. Adenosine A2A Receptors Contribute to the Radial Migration of Cortical Projection Neurons through the Regulation of Neuronal Polarization and Axon Formation.Cerebral Cortex. 31. (12). 5652–5663. 2021. 10.1093/cercor/bhab188.

[58] Domenici, M. R., Ferrante, A., Martire, A., Chiodi, V., Pepponi, R., Tebano, M. T., Popoli, P. Adenosine A2A receptor as potential therapeutic target in neuropsychiatric disorders.Pharmacological Research. 147. 104338. 2019. 10.1016/j.phrs.2019.104338.

[59] López-Cruz, L., Carbó-Gas, M., Pardo, M., Bayarri, P., Valverde, O., Ledent, C., Salamone, J. D., Correa, M. Adenosine A 2A receptor deletion affects social behaviors and anxiety in mice: Involvement of anterior cingulate cortex and amygdala.Behavioural Brain Research. 321. 8–17. 2017. 10.1016/j.bbr.2016.12.020.

[60] Eguchi, K., Velicky, P., Hollergschwandtner, E., Itakura, M., Fukazawa, Y., Danzl, J. G., Shigemoto, R. Advantages of Acute Brain Slices Prepared at Physiological Temperature in the Characterization of Synaptic Functions.Frontiers in Cellular Neuroscience. 14. 63. 2020. 10.3389/fncel.2020.00063.

[61] Fox, W. M. Reflex-ontogeny and behavioural development of the mouse.Animal Behaviour. 13. (2). 234–241. 1965. 10.1016/0003-3472(65)90041-2.

[62] Feather-Schussler, D. N., Ferguson, T. S. A Battery of Motor Tests in a Neonatal Mouse Model of Cerebral Palsy.Journal of Visualized Experiments: JoVE. (117). 53569. 2016. 10.3791/53569.

[63] Vignon, A. N., Dudon, G., Oliva, G., Thirard, S., Alenda, U. G., Brugoux, A., Cazevieille, C., Imbert, J., Bellières, C., Lehmann, S., Crozet, C., Torrent, J., Bertaso, F., Le Merrer, J., Becker, J. A. J., Perrier, V. Lifelong exposure to polystyrene-nanoplastics induces an attention-deficit hyperactivity disorder-like phenotype and impairs brain aging in mice.Journal of Hazardous Materials. 494. 138640. 2025. 10.1016/j.jhazmat.2025.138640.

[64] Jones, G., Willett, P., Glen, R. C., Leach, A. R., Taylor, R. Development and validation of a genetic algorithm for flexible docking.Journal of Molecular Biology. 267. (3). 727–748. 1997. 10.1006/jmbi.1996.0897.

[65] Derieux, C., Léauté, A., Brugoux, A., Jaccaz, D., Terrier, C., Pin, J.-P., Kniazeff, J., Le Merrer, J., Becker, J. A. J. Chronic sodium bromide treatment relieves autistic-like behavioral deficits in three mouse models of autism.Neuropsychopharmacology: Official Publication of the American College of Neuropsychopharmacology. 47. (9). 1680–1692. 2022. 10.1038/s41386-022-01317-1.

[66] Le Merrer, J., Detraux, B., Gandía, J., De Groote, A., Fonteneau, M., Kerchove d’Exaerde, A. de, Becker, J. A. J. Balance Between Projecting Neuronal Populations of the Nucleus Accumbens Controls Social Behavior in Mice.Biological Psychiatry. 95. (2). 123–135. 2024. 10.1016/j.biopsych.2023.05.008.

