## Supplementary Figures and Tables for "Neurobiological alterations and autism-like phenotype in mice following regulatory-limit exposure to fungicides in drinking water"

**Supplementary Figure S1**. DCX, NeuN, and ALDH1L1, P/T-Akt and P/T-β-catenin immunoblots with β-actin loading controls corresponding to immunoblots Figure 4 panels a,b,c, e,f.

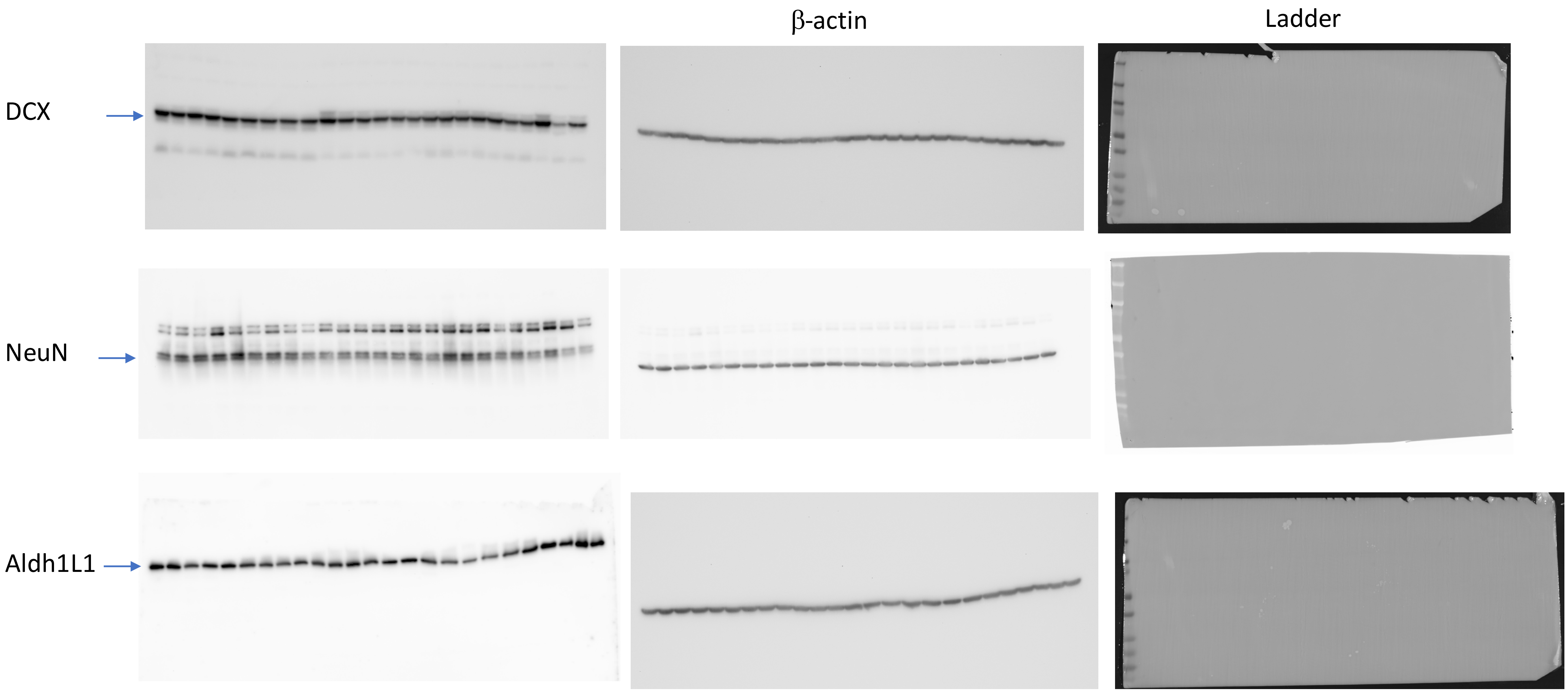

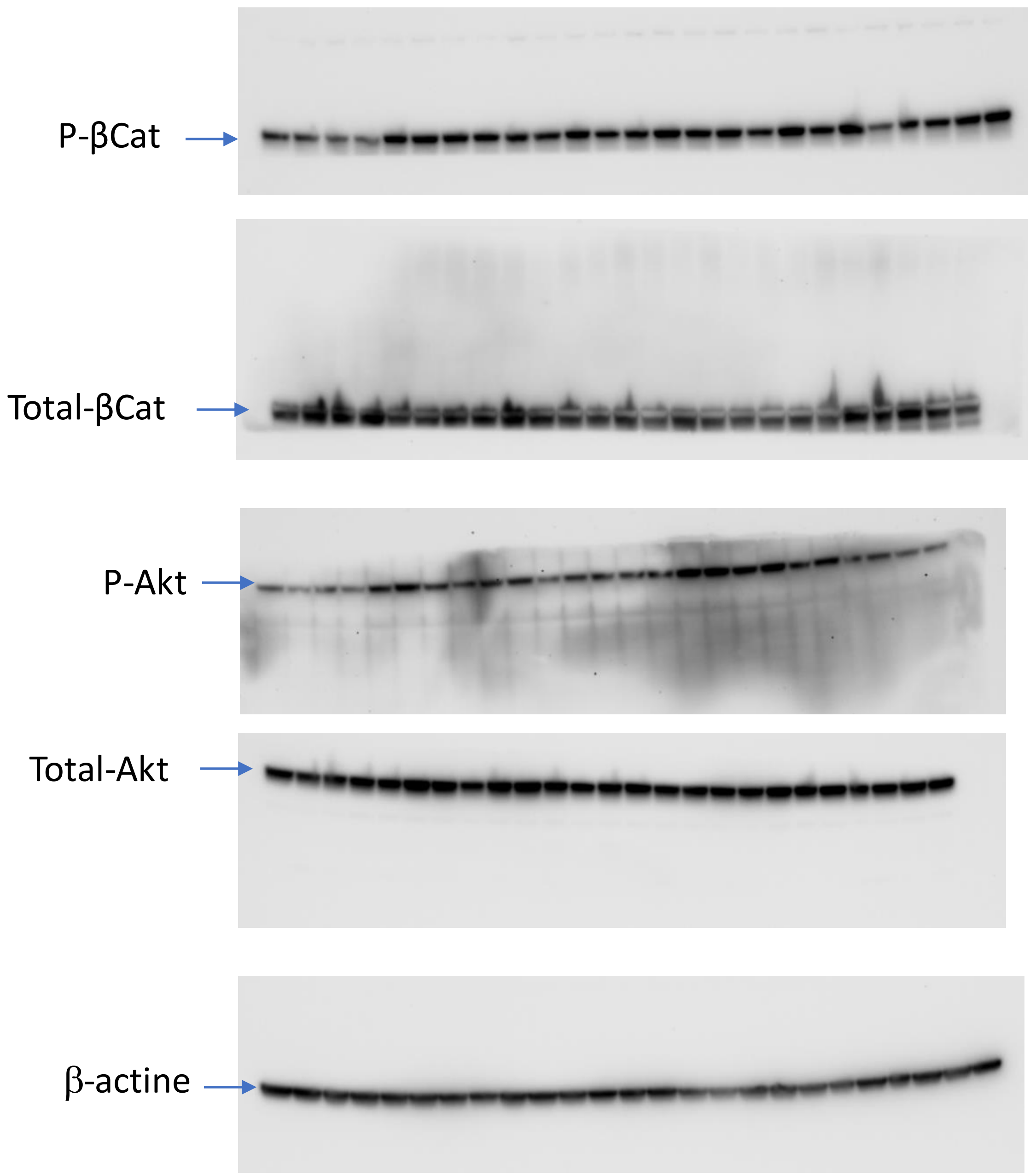


Ladder β-catenin

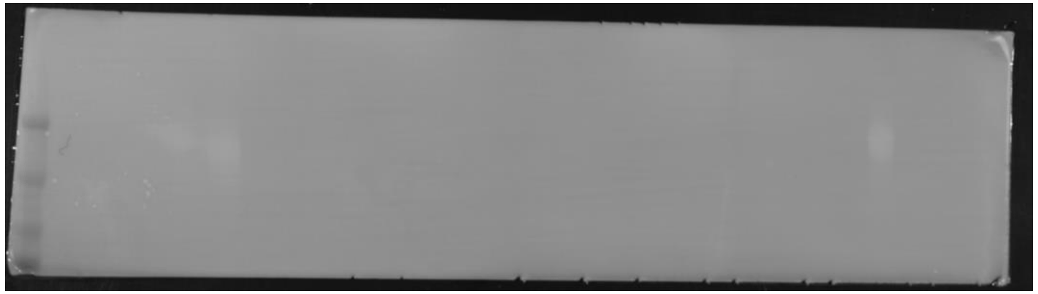

Ladder Akt

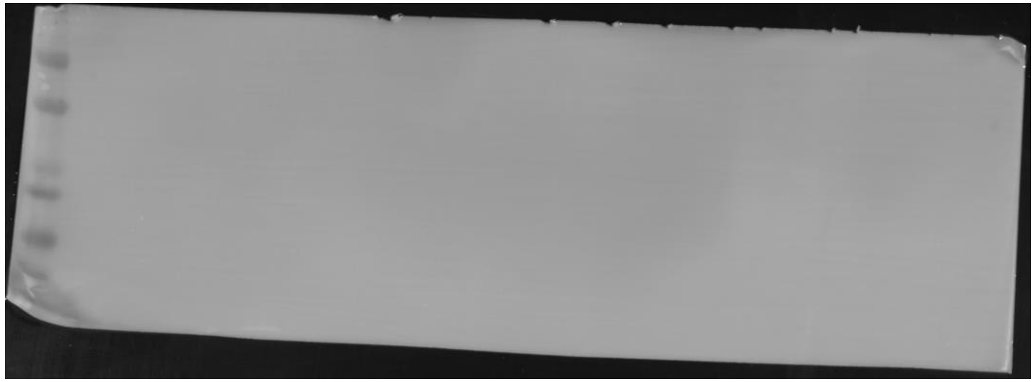

Same membrane : revelation with β-actin, then membrane was cut followed by probing with P-βCat/total-βCat and P-Akt/Total-Akt.

**Supplementary Figure S2**. P/T CAMKII and PSD95 immunoblots with β-actin loading controls corresponding to immunoblots Figure 5b.

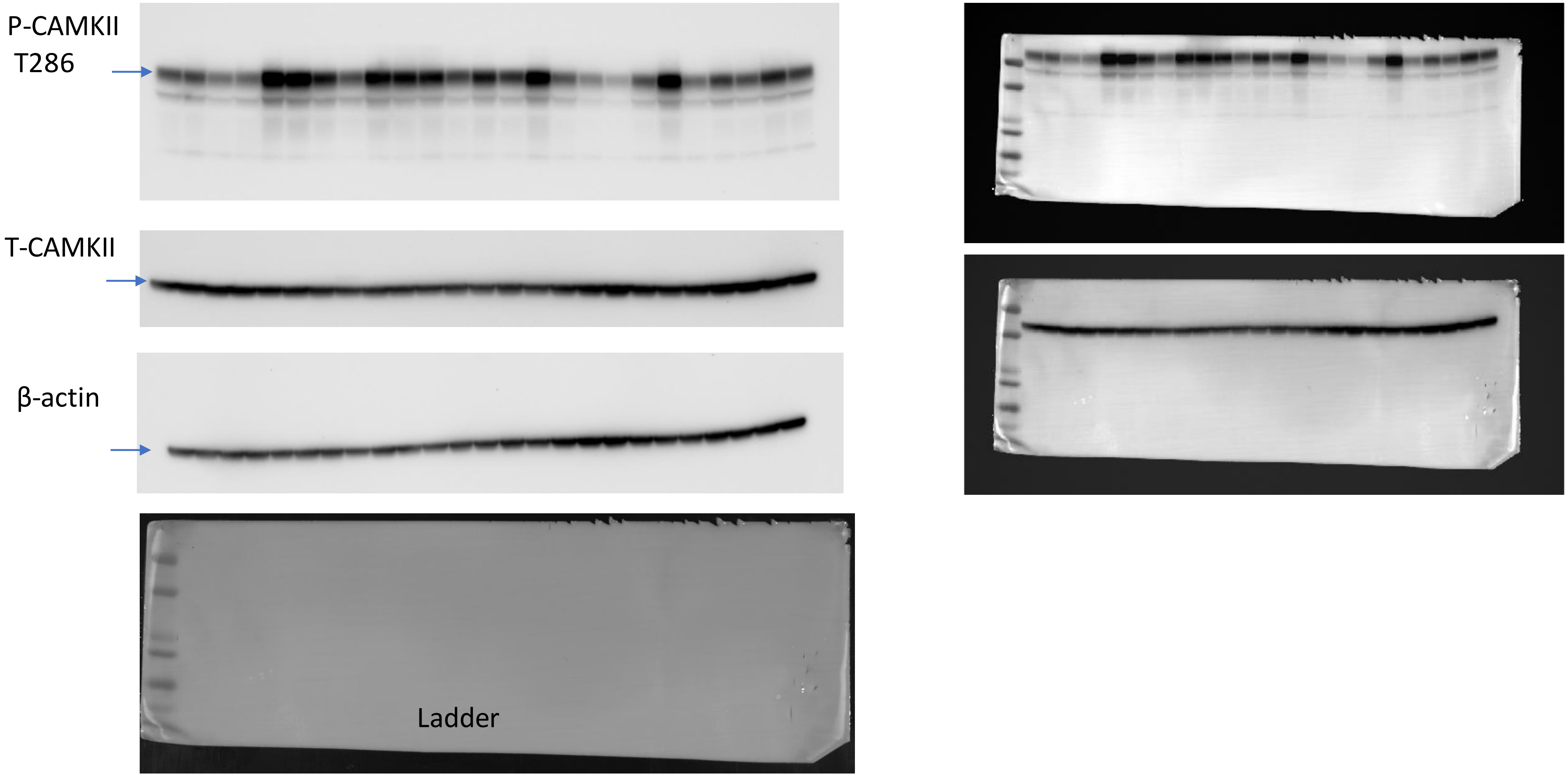

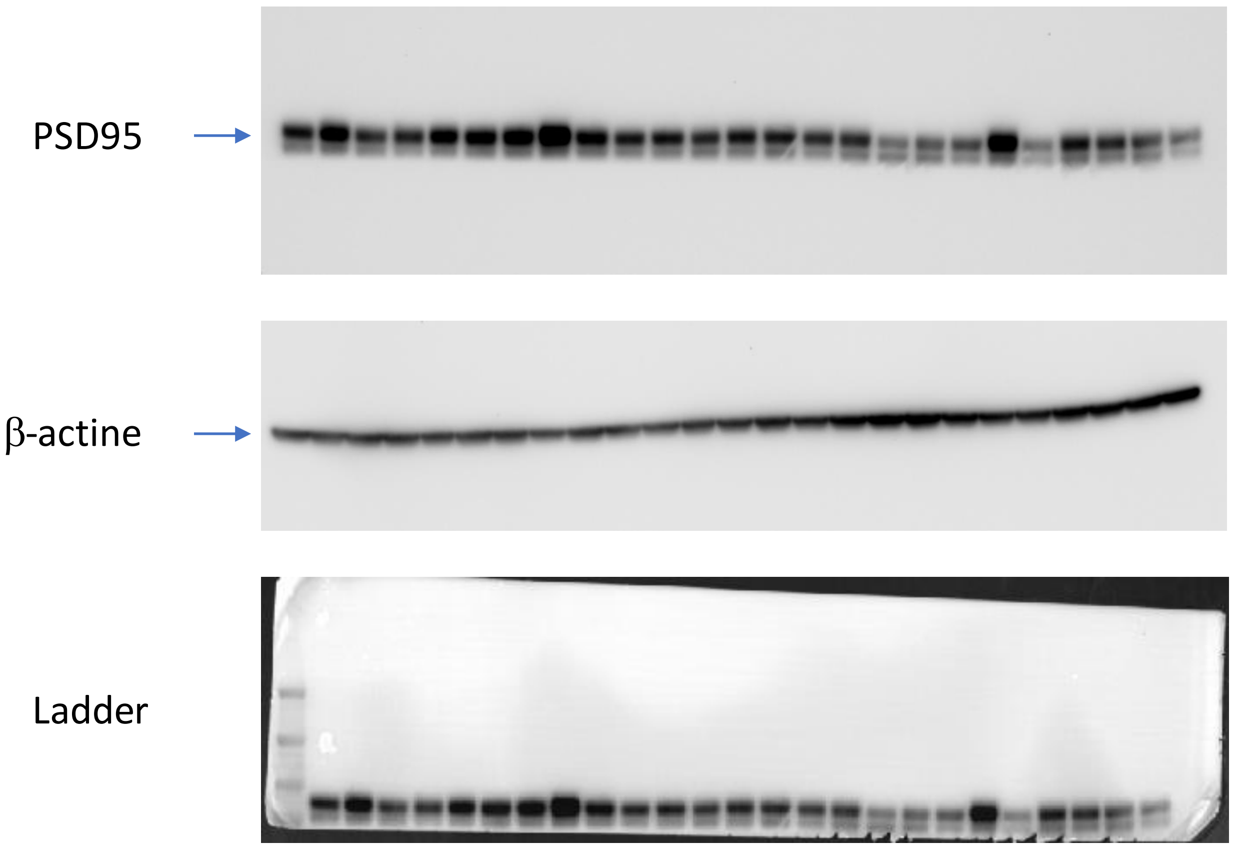


Membrane was stripped to remove P-CAMKII/T-CAMKII and further probe with PSD95 primary antibody
Same β-actin for the two immunoblots

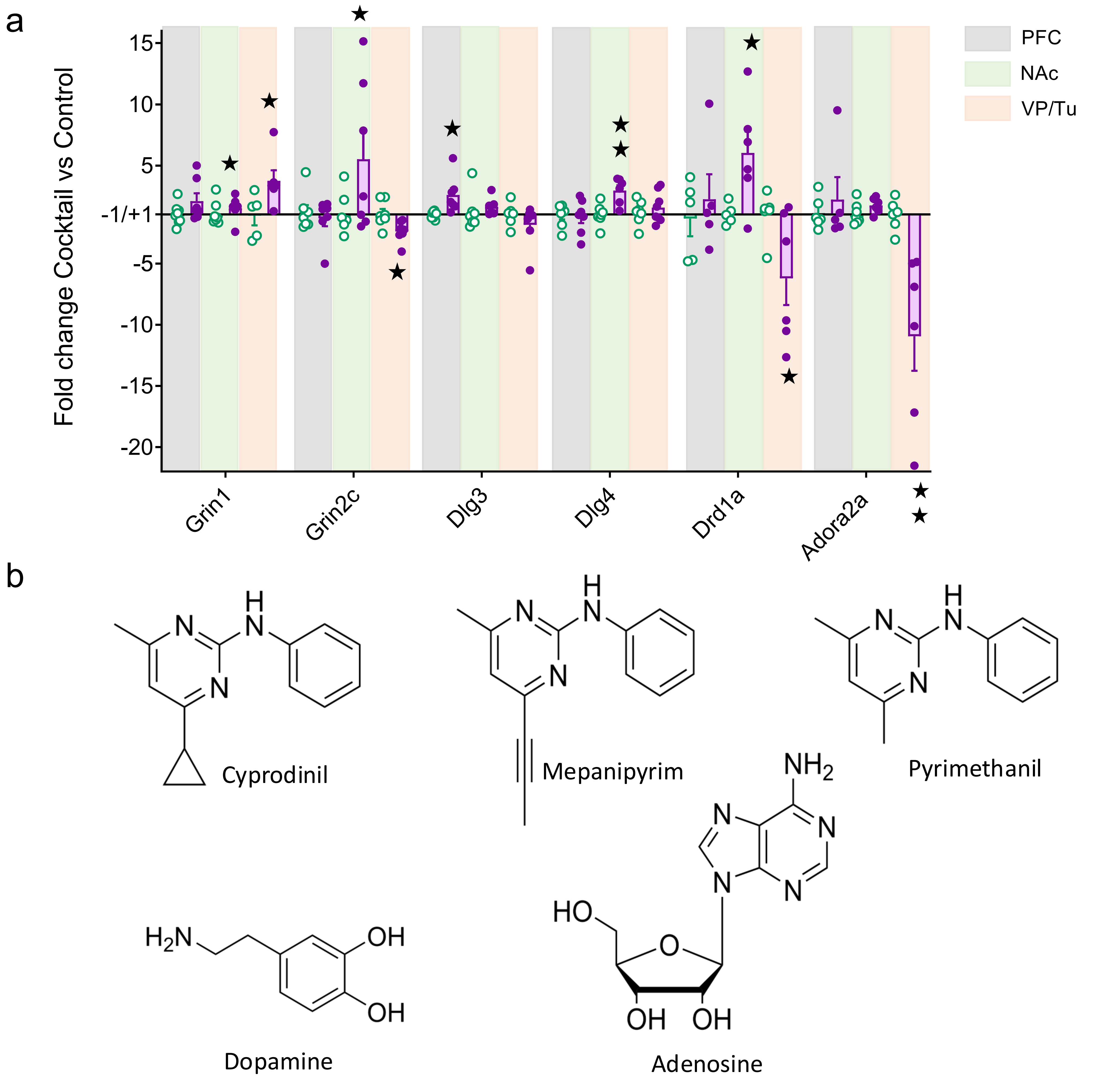


**Supplementary Figure S3**. **Transcriptional changes of fungicide-exposure in different brain regions.** The expression of 27 genes of interest coding for key players of glutamatergic neurotransmission and marker genes of striatal projection neurons (SPNs) was assessed at 3-month age in the prefrontal cortex (PFC), nucleus accumbens (NAc) and ventral pallidum/olfactory tuber (VP/Tu) of previously phenotyped control and fungicide-exposed mice (CTR n = 8, Cocktail n = 8). **(a)** All-life fungicide-exposure modified the expression of several genes Grin1 and Grin2c (coding for the NR1 and NR2C subunits of the NMDA receptor, respectively), *Dlg3* and *Dlg4* (post-synaptic proteins PSD93 and PSD95, respectively) and Adora2a (coding for the adenosine A2A receptor). **(b)** Chemical structures of anilinopyrimidine fungicides and endogenous ligands of adenosine and dopamine receptors.

**a**

Active form $A_{2A}$ (5G53)

| Ligand | Docking scores (a.u.) | p stacking |
|---|---|---|
| Adenosine (ADO) substrate | 65.36 | - |
| NECA agonist | 72.04 | 1 (168A) |
| ZM241385 antagonist | 71 | - |
| Cyprodinil | 58.79 | 1 (168A) |
| Mepanipyrim | 58.2 | 1 (168A) |
| Pyrimethanil | 51.81 | 1 (168A) |

**b**

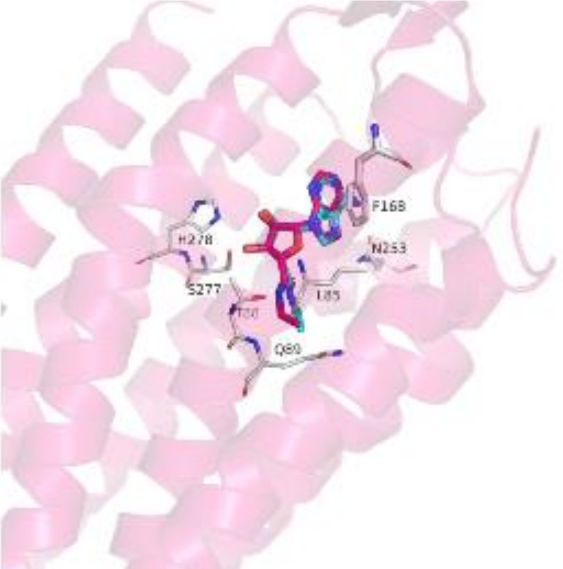


**c** **d** **e**

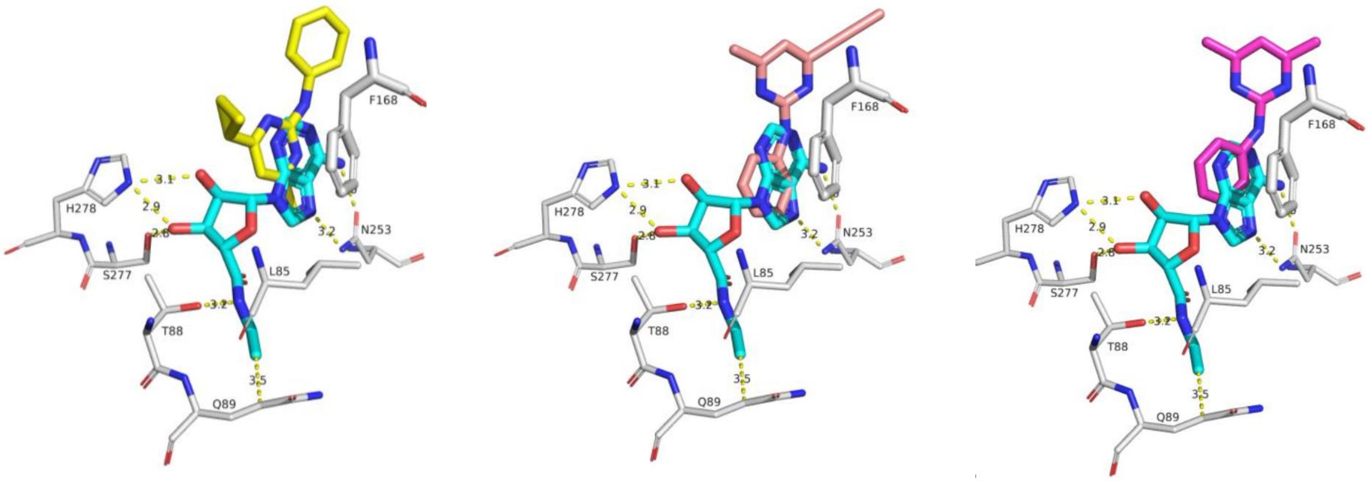


**Supplementary Figure S4 : Binding predictions by molecular docking of anilinopyrimidine fungicides in the active $A_{2A}$ cryo-EM structure.** (**a**) Docking scores and p stacking in the active form of $A_{2A}$ receptor (PDB 5G53) for control ligands (adenosine: ADO; agonist: NECA; antagonist: ZM241341) compared with anilinopyrimidine fungicides. (**b**) Overview of the docking on A2a receptor and control with NECA form crystal (cyan sticks) and docking pose (red sticks). Zoom-in views of docking poses show the NECA ligand (cyan) overlaid with (**c**) cyprodinil (yellow), (**d**) mepanipyrim (pale rose) and (**e**) pyrimethanil (pink sticks). Key residues of the $A_{2A}$ receptor are represented as thin green sticks, excepted F168, which is shown in thicker sticks.

Table S1. List of antibodies used for western-blot

| Antibody | Host | Clone | Supplier | Catalog No. | RRID | Application | Dilution | Target |
|---|---|---|---|---|---|---|---|---|
| Anti-Doublecortin (DCX) | Rabbit | Polyclonal | Cell Signaling Technology | 4604S | AB_10693771 | WB | 1:1000 | Doublecortin (DCX) |
| Anti-NeuN | Mouse | A60 | Merck Millipore (Chemicon) | MAB377 | AB_2298772 | WB | 1:1000 | NeuN (RBFOX3) |
| Anti-phospho-CaMKII (Thr286) | Rabbit | Polyclonal | Abcam | ab32678 | AB_731785 | WB | 1:1000 | Phospho-CaMKII (Thr286) |
| Anti-CaMKIIα | Mouse | 6G9 | Abcam | ab22609 | AB_447192 | WB | 1:1000 | CaMKIIα (CAMK2A) |
| Anti-NMDAR1 (GluN1) | Rabbit | EPR2481(2) | Abcam | ab109182 | AB_10861978 | WB | 1:5000 | NMDAR1 (GRIN1) |
| Anti-ALDH1L1 | Rabbit | Polyclonal | Abcam | ab87117 | AB_10712968 | WB | 1:1000 | ALDH1L1 |
| Anti-PSD95 (DLG4) | Rabbit | D27E11 | Cell Signaling Technology | 3450S | AB_2292883 | WB | 1:1000 | PSD95 (DLG4) |
| Anti-phospho-Akt (Ser473) | Rabbit | Polyclonal | Cell Signaling Technology | 9271S | AB_329825 | WB | 1:1000 | Phospho-Akt (Ser473; AKT1/2/3) |
| Anti-Akt (pan) | Rabbit | Polyclonal | Cell Signaling Technology | 9272S | AB_329827 | WB | 1:1000 | Total Akt (AKT1/2/3) |
| Anti-phospho-GSK-3β (Ser9) | Rabbit | Polyclonal | Cell Signaling Technology | 9336S | — | WB | 1:1000 | Phospho-GSK-3β (Ser9) |
| Anti-GSK-3β | Mouse | 7/GSK-3β | BD Biosciences | 610202 | AB_397601 | WB | 1:10000 | GSK-3β (GSK3B) |
| Anti-phospho-β-Catenin (Ser552) | Rabbit | Polyclonal | Cell Signaling Technology | 9966S | AB_331149 | WB | 1:1000 | Phospho-β-Catenin (Ser552; CTNNB1) |
| Anti-β-Catenin | Mouse | 14/Beta-Catenin | BD Biosciences | 610154 | AB_397555 | WB | 1:2000 | β-Catenin (CTNNB1) |
| Anti-β-Actin | Mouse | AC-15 | Sigma-Aldrich (Merck) | A5441 | AB_476744 | WB | 1:2000 | β-Actin (ACTB) |

**Table S2. List of primers used for qRT-PCR**

| Ref seq | Gene name | Gene title | Forward oligonucleotide | Reverse oligonucleotide |
|---|---|---|---|---|
| NM_009630 | adenosine A2a receptor | *Adora2a* | TCAGCCTCTTGGCTATTGCC | CTCAAACAGACAGGTCACCCG |
| NM_013464 | aryl-hydrocarbon receptor | *Ahr* | AGCACCACTAGACTGAGGGAT | TTCAGGTGCTGTAAGAAACACC |
| NM_019467 | allograft inflammatory factor 1 (Iba-1) | *Aif1* | TTGGAGGTGGATGTCACACG | TCTGACTCTGGCTCACGACT |
| NM_001177780 | discs, large homolog 3 (Drosophila) | *Dlg3* | AAGGCAGCAGCTTTCTCTTG | AATCAACACTTCCCGCTCAC |
| NM_007864 | discs, large homolog 4 (Drosophila) | *Dlg4* | AAGCTGGAGCAGGAGTTCAC | GAGGTCTTCGATGACACGTT |
| NM_010076 | dopamine receptor D1A | *Drd1a* | AGATCGGGCATTTGGAGAG | GGATGCTGCCTCTTCTTCTG |
| NM_010077 | dopamine receptor D2 | *Drd2* | TGCCATTGTTCTTGGTGTGT | GTGAAGGCGCTGTAGAGGAC |
| NM_001113325 | glutamate receptor, ionotropic, AMPA1 (alpha 1) | *Gria1* | AACGAGGGATGTGTGGAAAG | AGAGTTTTCCCACTGCTCCA |
| NM_013540 | glutamate receptor, ionotropic, AMPA2 (alpha 2) | *Gria2* | CGTGCTGATGTGGGAATATG | GGACCCAGGCTAGTTTCTCC |
| NM_016886 | glutamate receptor, ionotropic, AMPA3 (alpha 3) | *Gria3* | TGAAGAAGCGGGTAAATTGG | GCCAAACTCCCATAGCTTGA |
| NM_019691 | glutamate receptor, ionotropic, AMPA4 (alpha 4) | *Gria4* | TGGCACTCATCGTACTGCAT | TGCTGGGGATACAACTGTGA |
| NM_001177656 | glutamate receptor, ionotropic, NMDA1 (zeta 1) | *Grin1* | TAGGGCTCCAGACTCCAAGA | AAGGTTGGGTGAGTGACACC |
| NM_008170 | glutamate receptor, ionotropic, NMDA2A (epsilon 1) | *Grin2a* | GTCTGGAGGACAGCAAGAGG | CGAGGGACATCTCCCAATAA |
| NM_008171 | glutamate receptor, ionotropic, NMDA2B (epsilon 2) | *Grin2b* | AACGAAGCTGCCATGTCTCT | AGCAGTCGAGTATGGGGATG |
| NM_010350 | glutamate receptor, ionotropic, NMDA2C (epsilon 3) | *Grin2c* | CAGTGGGGTGCTAGAAGAGG | ACCAGCCACAGAATCCAGTC |
| NM_001160353 | glutamate receptor, metabotropic 2 | *Grm2* | CTTGTAGCTATGCCCCGTGT | GACTGGAAGCACCTTTGCAT |
| NM_001013385 | glutamate receptor, metabotropic 4 | *Grm4* | CTTTCTCTGCTATGCCACCACC | TAGCTGATGCTCATGCCAAGCC |
| NM_001143834 | glutamate receptor, metabotropic 5 | *Grm5* | CCACTGACGACTTGACAGGTT | CAGTAACGAAGAGGGTGGCTA |
| NM_031168 | interleukin 6 | *Il6* | AGTCCTTCCTACCCCAATTTCC | ATGAATTGGATGGTCTTGGTCCT |
| NM_011025 | oxytocin | *Oxt* | CTGCTTGGCTTACTGGCTCT | GGGAGACACTTGCGCATATC |
| NM_027571 | purinergic receptor P2Y, G-protein coupled 12 | *P2ry12* | TCCTTAACACTAGAGGCAGCA | TCAAGCAGCAGGCATTTTCA |
| NM_018863 | prodynorphin | *Pdyn* | TTTGGCAACGGAAAAGAATC | TAGCGTTTGGCCTGTTTTCT |
| NM_001002927 | preproenkephalin | *Penk* | ATGCAGATGAGGGAGACACC | GCTTCTGCAGCTCTTTTGCT |
| NM_009194 | solute carrier family 12, member 2 | *Slc12a2* | AGGTAAAACATCCGGTGGGT | AGCACAAAGAGAAAGACGCAAC |
| NM_020333 | solute carrier family 12, member 5 | *Slc12a5* | CAGACCTATGTGCAGGGCAA | CCGAGTCGGGATGCGAAATA |
| NM_010020 | solute carrier family 6 (neurotransmitter transporter, dopamine), member 3 | *Slc6a3* | TGAACTTTGGCCTAGCCTGT | TCAAGATCTTCGTGCCACTG |
| NM_010484 | solute carrier family 6 (neurotransmitter transporter, serotonin), member 4 | *Slc6a4* | CCCTCTAAGCCAAGCTGATG | CCGTGTCACCAGCTAATGTG |

**Table S3**: Transcription levels of a set of 27 genes in the PFC, Nac and VP in fongicide exposed animal vs contrôle

| | | | PFC | | | | NAc | | | | VP | | | |
|---|---|---|---|---|---|---|---|---|---|---|---|---|---|---|
| Refseq | Gene name | Gen title | FC | sem | p values | Benjamini-Hochberg Adjusted P value | FC | sem | p values | Benjamini-Hochberg Adjusted P value | FC | sem | p values | Benjamini-Hochberg Adjusted P value |
| NM_009630 | adenosine A2a receptor | *Adora2a* | 2.22 | 1.45 | 0.543 | 0.738 | 1.63 | 0.29 | 0.249 | 0.518 | **-10.93** | **2.44** | **0.006** | 0.139 |
| NM_013464 | aryl-hydrocarbon receptor | *Ahr* | **2.45** | **0.54** | **0.032** | 0.123 | -1.03 | 0.37 | 0.917 | 0.918 | **3.597** | **0.47** | **0.024** | 0.168 |
| NM_019467 | allograft inflammatory factor 1 (Iba-1) | *Aif1* | -1.88 | 1.37 | 0.553 | 0.738 | **3.01** | **0.53** | **0.015** | 0.102 | 3.95 | 1.29 | 0.171 | 0.462 |
| NM_001177780 | discs, large homolog 3 (Drosophila) | *Dlg3* | 2.59 | 0.52 | 0.015 | 0.123 | 1.63 | 0.26 | 0.516 | 0.750 | -1.83 | 0.60 | 0.320 | 0.617 |
| NM_007864 | discs, large homolog 4 (Drosophila) | *Dlg4* | -1.18 | 0.49 | 0.871 | 0.904 | **2.95** | **0.38** | **0.003** | 0.099 | 1.56 | 0.43 | 0.507 | 0.618 |
| NM_010076 | dopamine receptor D1A | *Drd1a* | 2.24 | 1.61 | 0.544 | 0.738 | **6.02** | **1.52** | **0.032** | 0.150 | **-6.21** | **1.89** | **0.047** | 0.212 |
| NM_010077 | dopamine receptor D2 | *Drd2* | 6.01 | 2.58 | 0.136 | 0.279 | 1.39 | 0.17 | 0.918 | 0.918 | -1.44 | 1.34 | 0.522 | 0.618 |
| NM_001113325 | glutamate receptor, ionotropic, AMPA1 (alpha 1) | *Gria1* | 2.12 | 0.55 | 0.085 | 0.229 | 1.36 | 0.16 | 0.347 | 0.612 | 1.45 | 0.34 | 0.307 | 0.617 |
| NM_013540 | glutamate receptor, ionotropic, AMPA2 (alpha 2) | *Gria2* | 2.5 | 0.89 | 0.128 | 0.279 | 2.50 | 0.91 | 0.165 | 0.381 | **-3.21** | **0.50** | **0.010** | 0.139 |
| NM_016886 | glutamate receptor, ionotropic, AMPA3 (alpha 3) | *Gria3* | 1.65 | 0.81 | 0.637 | 0.781 | -1.21 | 0.29 | 0.581 | 0.750 | 1.46 | 0.14 | 0.282 | 0.617 |
| NM_019691 | glutamate receptor, ionotropic, AMPA4 (alpha 4) | *Gria4* | **1.62** | **0.2** | **0.037** | 0.123 | **1.46** | **0.11** | **0.039** | 0.150 | 1.76 | 0.17 | 0.427 | 0.618 |
| NM_001177656 | glutamate receptor, ionotropic, NMDA1 (zeta 1) | *Grin1* | 2.08 | 0.59 | 0.166 | 0.298 | 1.49 | 0.36 | 0.493 | 0.750 | **3.74** | **0.75** | **0.041** | 0.212 |
| NM_008170 | glutamate receptor, ionotropic, NMDA2A (epsilon 1) | *Grin2a* | 1.49 | 0.92 | 0.76 | 0.855 | 1.05 | 0.18 | 0.845 | 0.913 | 1.62 | 0.95 | 0.624 | 0.703 |
| NM_008171 | glutamate receptor, ionotropic, NMDA2B (epsilon 2) | *Grin2b* | 1.78 | 0.53 | 0.292 | 0.463 | 1.14 | 0.23 | 0.592 | 0.750 | 1.00 | 0.37 | 0.851 | 0.883 |
| NM_010350 | glutamate receptor, ionotropic, NMDA2C (epsilon 3) | *Grin2c* | -1.31 | 0.6 | 0.574 | 0.738 | 5.51 | 2.15 | 0.121 | 0.326 | **-2.37** | **0.30** | **0.017** | 0.156 |
| NM_001160353 | glutamate receptor, metabotropic 2 | *Grm2* | -1.39 | 1.49 | 0.805 | 0.869 | 1.86 | 0.49 | 0.169 | 0.381 | 1.82 | 0.43 | 0.657 | 0.709 |
| NM_001013385 | glutamate receptor, metabotropic 4 | *Grm4* | 1.31 | 0.62 | 0.668 | 0.784 | 2.24 | 0.24 | 0.012 | 0.102 | 1.99 | 0.89 | 0.445 | 0.618 |
| NM_001143834 | glutamate receptor, metabotropic 5 | *Grm5* | **3.23** | **0.82** | **0.041** | 0.123 | 1.17 | 0.20 | 0.747 | 0.840 | -1.63 | 0.95 | 0.498 | 0.618 |
| NM_031168 | interleukin 6 | *Il6* | **2.46** | **0.34** | **0.012** | 0.123 | 2.47 | 0.57 | 0.096 | 0.290 | 1.84 | 0.43 | 0.455 | 0.618 |
| NM_011025 | oxytocin | *Oxt* | 3.78 | 1.2 | 0.103 | 0.252 | **3.65** | **0.50** | **0.009** | 0.102 | 1.57 | 0.70 | 0.891 | 0.891 |
| NM_027571 | purinergic receptor P2Y, G-protein coupled 12 | *P2ry12* | 2.97 | 1.03 | 0.145 | 0.279 | -1.34 | 0.22 | 0.611 | 0.750 | -2.66 | 1.31 | 0.432 | 0.618 |
| NM_018863 | prodynorphin | *Pdyn* | 3.38 | 0.89 | 0.036 | 0.123 | 1.66 | 0.58 | 0.362 | 0.612 | 2.97 | 0.72 | 0.119 | 0.359 |
| NM_001002927 | preproenkephalin | *Penk* | 1.78 | 0.43 | 0.203 | 0.342 | **3.18** | **0.61** | **0.038** | 0.150 | 2.75 | 0.73 | 0.461 | 0.618 |
| NM_009194 | solute carrier family 12, member 2 | *Slc12a2* | **2.16** | **0.26** | **0.009** | 0.123 | 1.08 | 0.12 | 0.663 | 0.778 | -2.03 | 0.51 | 0.226 | 0.554 |
| NM_020333 | solute carrier family 12, member 5 | *Slc12a5* | -1.00 | 0.68 | 0.944 | 0.944 | 1.21 | 0.26 | 0.534 | 0.750 | 2.84 | 0.86 | 0.082 | 0.278 |
| NM_010020 | solute carrier family 6 (neurotransmitter transporter, dopamine), member 3 | *Slc6a3* | **5.04** | **1.21** | **0.025** | 0.123 | 2.18 | 0.97 | 0.294 | 0.567 | -2.02 | 1.44 | 0.526 | 0.618 |
| NM_010484 | solute carrier family 6 (neurotransmitter transporter, serotonin), member 4 | *Slc6a4* | 4.54 | 0.65 | 0.039 | 0.123 | 3.84 | 1.22 | 0.095 | 0.290 | -12.51 | 4.29 | 0.069 | 0.267 |

Table S4. Number of mice analyzed by multi-electrode array (MEA) and incidence of epileptiform activity

| Postnatal age | Control | Cocktail |
|---|---|---|
| **P10** | 0/5 (0%) | 3/5 (60%) |
| **P18** | 0/5 (0%) | 4/5 (80%) |
| **P30** | 1/5 (20%) | 2/5 (40%) |
